# *In vitro* Lung Epithelial Cell Model Reveals Novel Roles for *Pseudomonas aeruginosa* Siderophores

**DOI:** 10.1101/2023.01.26.525796

**Authors:** Donghoon Kang, Qi Xu, Natalia V. Kirienko

**Affiliations:** Department of BioSciences, Rice University, Houston, TX, USA; Department of Bioengineering, Rice University, Houston, TX, USA

**Author notes:** **Corresponding Author:** Natalia V. Kirienko.

**Keywords:** *Pseudomonas aeruginosa*, Virulence, Siderophores, Lung Epithelial Cells, Inflammation, Pyoverdine, Pyochelin, Rhamnolipids

## Abstract

Multidrug-resistant *Pseudomonas aeruginosa* is a common nosocomial respiratory pathogen that continues to threaten the lives of patients with mechanical ventilation in intensive care units and those with underlying comorbidities such as cystic fibrosis or chronic obstructive pulmonary disease. For over 20 years, studies have repeatedly demonstrated that the major siderophore pyoverdine is an important virulence factor for *P. aeruginosa* in invertebrate and mammalian hosts *in vivo*. Despite its physiological significance, an *in vitro,* mammalian cell culture model to characterize the impact and molecular mechanism of pyoverdine-mediated virulence has only been developed very recently. In this study, we adapt a previously-established, murine macrophage-based model for human bronchial epithelial cells (16HBE). We demonstrate that conditioned medium from *P. aeruginosa* induced rapid 16HBE cell death through the pyoverdine-dependent secretion of cytotoxic rhamnolipids. Genetic or chemical disruption of pyoverdine biosynthesis decreased rhamnolipid production and mitigated cell death. Consistent with these observations, chemical depletion of lipid factors or genetic disruption of rhamnolipid biosynthesis was sufficient to abrogate conditioned medium toxicity. Furthermore, we also examine the effects of purified pyoverdine exposure on 16HBE cells. While pyoverdine accumulated within cells, the siderophore was largely sequestered within early endosomes, showing minimal cytotoxicity. More membrane-permeable iron chelators, such as the siderophore pyochelin, decreased epithelial cell viability and upregulated several proinflammatory genes. However, pyoverdine potentiated these iron chelators in activating proinflammatory pathways. Altogether, these findings suggest that the siderophores pyoverdine and pyochelin play distinct roles in virulence during acute *P. aeruginosa* lung infection.

## Introduction

Multidrug-resistant *Pseudomonas aeruginosa* is one of the most common Gram-negative, respiratory pathogens, and infects mechanically-ventilated patients in intensive care units or those with cystic fibrosis (CF) or chronic obstructive pulmonary disease (COPD) (1–5). This pathogen’s intrinsic resistance to several classes of antibiotics and exceptional ability to form biofilms on medical devices and airway tissue pose a serious challenge for medical intervention (6, 7). In addition to colonizing the respiratory tract, *P. aeruginosa* actively deploys numerous virulence factors and toxins that damage host tissue, affecting pulmonary function (8). Two of the major virulence factors produced by this pathogen are the siderophores pyoverdine and pyochelin.

Several have proposed possible mechanisms of siderophore-dependent virulence during *P. aeruginosa* lung infection (9–12). As siderophores, both pyoverdine and pyochelin scavenge ferric iron and provide the pathogen with this essential micronutrient during infection. However, pyoverdine exhibits orders of magnitude higher affinity for ferric iron and is distinctly able to chelate the metal from host ferroproteins such as transferrin and lactoferrin (13, 14). Generally, iron acquisition serves an important function during infection by promoting bacterial growth and biofilm formation (15, 16), and *P. aeruginosa* mutants lacking various iron uptake systems exhibit attenuation of virulence during murine lung infection (17). It is important to note however that these iron uptake systems do not contribute equally, and of the two siderophores, pyoverdine plays a greater role in *P. aeruginosa* virulence (17).

Pyoverdine-mediated iron uptake further promotes *P. aeruginosa* virulence by derepressing the alternative sigma factor PvdS, which activates the transcription of several virulence genes such as those encoding the translational inhibitor exotoxin A, the exoprotease PrpL (protease IV), and pyoverdine biosynthetic enzymes (12, 18). Furthermore, we have recently used a *Caenorhabditis elegans* nematode model to demonstrate that pyoverdine likely directly chelates host iron, disrupting mitochondrial homeostasis (19–21). Pyoverdine’s well documented role in acute lung infection is likely mediated by a combination of these various pathogenic functions (17, 22–25).

Recently, we established the first-reported *in vitro* cell culture model for pyoverdine-dependent virulence, where murine macrophages were treated with conditioned medium from *P. aeruginosa* grown in serum-free cell culture medium (26). Under these conditions, *P. aeruginosa* exhibited robust pyoverdine production, yet the siderophore was not required for bacterial growth (**Fig. 1A, B; Fig. S1A, B**), allowing for the study of pyoverdine’s role in virulence. This pyoverdine-rich conditioned medium from wild-type *P. aeruginosa* PAO1 was cytotoxic towards murine macrophages, including murine alveolar macrophages (**Fig. S1C)**; in clinical isolates; pyoverdine content in the conditioned medium positively correlated with cytotoxicity (26).

**Fig. 1.**
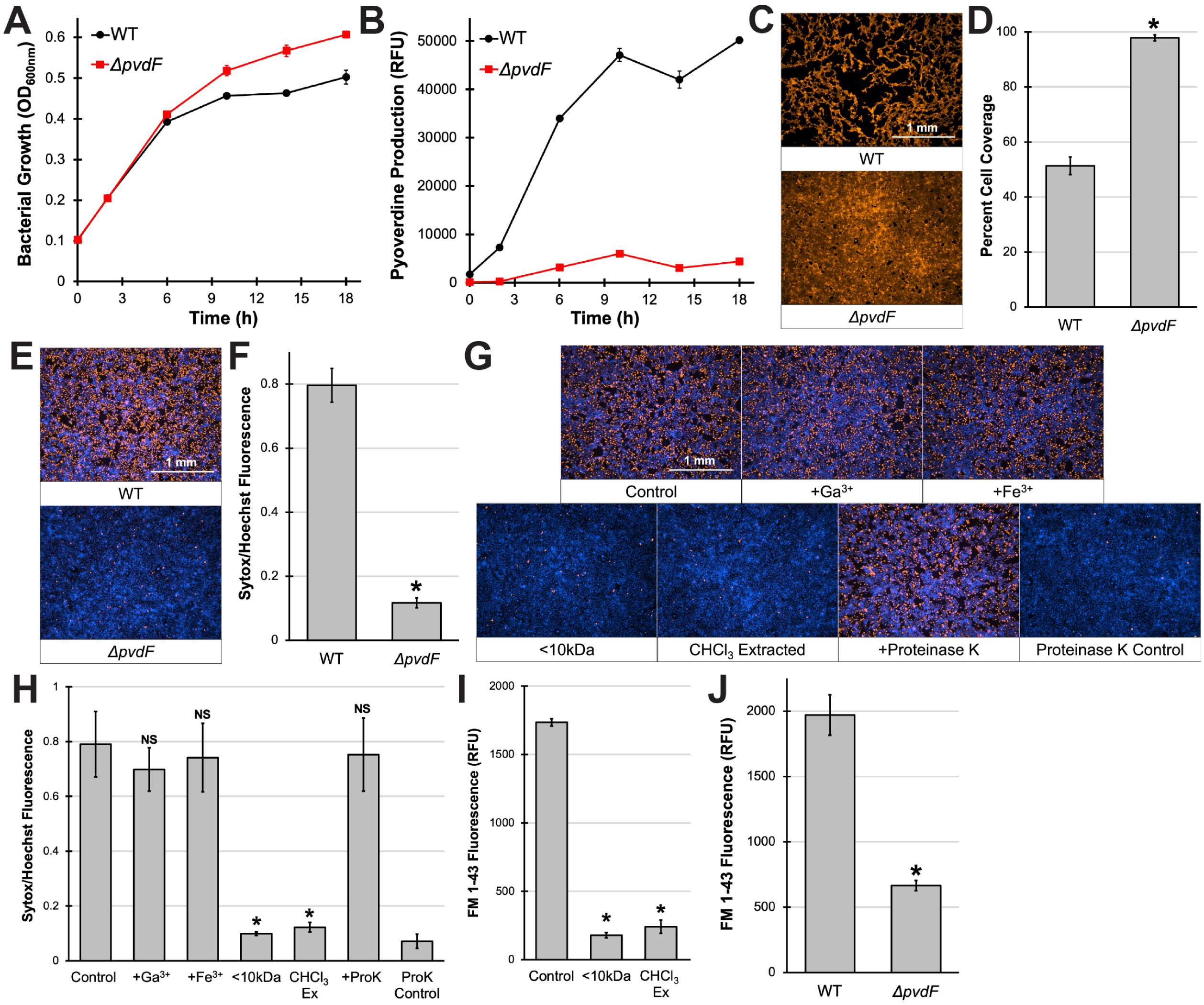
Pyoverdine-rich conditioned medium kills 16HBE cells and damages the epithelial monolayer. **(A, B)** Bacterial growth **(A)** or pyoverdine production **(B)** by WT *P. aeruginosa* PAO1 or pyoverdine biosynthetic mutant (PAO1*ΔpvdF*) in serum-free EMEM. **(C)** Fluorescent micrographs of 16HBE human bronchial epithelial cells after 30 min exposure to conditioned medium from WT PAO1 or PAO1*ΔpvdF* grown in EMEM. Cells were prelabeled with CellMask Orange plasma membrane stain. **(D)** Quantification of percentage micrograph area covered by fluorescent cells. **(E)** Fluorescent micrographs of 16HBE cells after 15 min exposure to conditioned medium from WT PAO1 or PAO1*ΔpvdF* in the presence of Sytox Orange nucleic acid stain (red). Cells were prelabeled with Hoechst 33342 nucleic acid stain (blue). **(F)** Quantification of Sytox Orange mean fluorescence intensity normalized to that of Hoechst 33342. **(G)** Fluorescent micrographs of 16HBE cells after 15 min exposure to conditioned medium from WT PAO1 in the presence of Sytox Orange nucleic acid stain (red). Conditioned medium was pretreated with 200 µM Ga(NO_3_)_3_, 200 µM FeCl_3_, or 100 µg/mL proteinase K for 24 h, had macromolecules depleted via a 10 kDa centrifugal filter, or had lipids depleted by chloroform (CHCl_3_) extraction. Cells were prelabeled with Hoechst 33342 nucleic acid stain (blue). **(H)** Quantification of Sytox Orange mean fluorescence intensity normalized to that of Hoechst 33342. **(I)** Quantification of lipids by FM 1-43 fluorescent labeling in conditioned medium from WT PAO1 (control) or macromolecule-(<10 kDa) or lipid-(CHCl_3_) depleted material. **(J)** Quantification of lipids in conditioned medium from WT PAO1 or PAO1*ΔpvdF* by FM 1-43 labeling. All error bars represent SEM from at least three biological replicates. * corresponds to *p* < 0.01 and NS corresponds to *p* > 0.05 based on Student’s *t*-test **(D, F, J)** or one-way ANOVA with Dunnett’s multiple comparisons test **(H, I)**.

In this report, we adapted this *in vitro* pyoverdine virulence model for human bronchial epithelial cells to examine the consequences of pyoverdine production (i.e. exposure to pyoverdine and/or pyoverdine-regulated virulence factors) during *P. aeruginosa* lung infection. Conditioned medium from *P. aeruginosa* caused acute cell death and severe damage to the epithelial monolayer in a pyoverdine-, but not pyochelin-, dependent manner. Interestingly, this damage did not require host iron chelation nor production of the two known pyoverdine-regulated toxins, exotoxin A or PrpL. Instead, pyoverdine production promoted the secretion of cytotoxic rhamnolipids that have previously been shown to permeabilize host membranes (27). Consistent with this observation, chemical depletion of lipids or genetic disruption of rhamnolipid production was sufficient to abrogate toxicity from conditioned medium on 16HBE cells. Importantly, the pyoverdine biosynthetic inhibitor 5-fluorocytosine effectively inhibited rhamnolipid production and mitigated *P. aeruginosa* virulence in two highly virulent clinical isolates. We also examined the effects of exposing 16HBE cells to purified pyoverdine. While pyoverdine accumulated within cells, the siderophore was largely sequestered within early endosomes, showing minimal cytotoxicity. More membrane-permeable iron chelators such as the siderophore pyochelin, decreased epithelial cell viability and upregulated several proinflammatory pathways. However, pyoverdine potentiated these iron chelators in activating proinflammatory pathways. Altogether, these findings suggest that the siderophores pyoverdine and pyochelin play distinct roles in virulence during acute *P. aeruginosa* lung infections.

## Results

### Pyoverdine-rich, conditioned medium induces rapid cell death and damages the epithelial monolayer

To investigate the role of pyoverdine production during *P. aeruginosa* lung infection, we treated human bronchial epithelial cells (16HBE) with bacteria-free, pyoverdine-rich conditioned medium from *P. aeruginosa* PAO1 grown in serum-free cell growth medium (Eagle’s Minimum Essential Medium or EMEM). To visualize the integrity of the epithelial monolayer, we prelabeled cells with a plasma membrane stain. Within 30 min, the conditioned medium severely damaged the monolayer, causing detachment of more than half of the cells **(Fig. 1C, D)**. This disruption was significantly attenuated in 16HBE cells treated with identically prepared material from an isogenic pyoverdine biosynthetic mutant (PAO1*ΔpvdF*) but not a pyochelin mutant (PAO1*ΔpchBA*) **(Fig. S1D, E)**. Preventing the biosynthesis of both pyoverdine and pyochelin (PAO1*ΔpvdFΔpchBA*) conferred no further protection to 16HBE cells than the disruption of pyoverdine alone (**Fig. S1D, E**). To determine whether cell detachment was caused by cell death (rather than from degradation of the extracellular matrix via bacterial proteases and other factors), we labeled cells with cell-permeant (Hoechst 33342; labels all cells) and cell-impermeant (Sytox Orange; labels only dead cells) nucleic acid stains. Exposure to pyoverdine-rich, conditioned medium from wild-type cells caused rapid (within 15 min) membrane permeabilization and internalization of the cell-impermeant nucleic acid stain, suggesting cell death **(Fig. 1E, F)**. In contrast, conditioned medium from the pyoverdine mutant exhibited substantially less cytotoxicity **(Fig. 1E, F)**.

One key difference between pyoverdine and pyochelin is their affinity for ferric iron. Due to an exceptionally high affinity for the metal, pyoverdine is uniquely able to remove iron from host ferroproteins (13, 14) and induce a lethal hypoxic response in a *C. elegans* nematode model (19). We thus examined whether host iron chelation was important for the cell death we observed in the epithelial monolayer. To hinder pyoverdine’s ability to bind iron, we pretreated the conditioned media with gallium (Ga^3+^) or ferric iron (Fe^3+^), either of which would prevent pyoverdine from scavenging iron from the epithelial cells. Surprisingly, even with the addition of excess metal, there was no significant rescue **(Fig. 1G, H; Fig. S2)**. Interestingly, we observed that removing material with a molecular mass greater than 10 kDa (via centrifugal filtration) prevented damage to the epithelial monolayer **(Fig. 1G, H, Fig. S2)**, suggesting that macromolecules or molecular complexes were responsible for conditioned medium cytotoxicity. To investigate the role of proteinaceous toxins, we pre-treated the conditioned medium with proteinase K. Degradation of secreted proteins did not significantly attenuate conditioned medium cytotoxicity **(Fig. 1G, H)**. We could not determine whether proteinase K treatment mitigated cell detachment; treatment alone caused considerable damage to the extracellular matrix, making it very difficult to unambiguously assess contributions **(Fig. S2)**. Consistent with this observation, genetic disruption of the two pyoverdine-regulated toxins, exotoxin A and PrpL, or the type II secretion system through which these toxins are secreted, did not affect conditioned medium toxicity **(Fig. S3)**. We did however observe that removing lipids from the conditioned medium by chloroform extraction abrogated cytotoxicity and prevented damage to the epithelial monolayer **(Fig. 1G, H; Fig. S2)**. Based on these results, we directly measured lipid content in the conditioned medium using the lipophilic dye FM 1-43. This probe is nonfluorescent in aqueous solution but is highly-fluorescent upon binding to lipid membranes (28). Chloroform extraction and removal of macromolecules (>10 kDa) significantly depleted lipid content **(Fig. 1I)**. Importantly, conditioned medium from the pyoverdine biosynthetic mutant also contained significantly less lipid material than that from wild-type PAO1 **(Fig. 1J)**.

### Pyoverdine regulates the production of cytotoxic rhamnolipids

Based on previous studies (27, 29), we posited that the relevant secreted lipid factors were rhamnolipids. We recently demonstrated that *P. aeruginosa* secretes rhamnolipids that rapidly induce membrane rupture and permeabilization in a wide range of host cells, including murine macrophages, human bronchial epithelial cells, and erythrocytes (27). Chloroform extraction and centrifugal filtration, two treatments that would likely remove these rhamnolipids from conditioned medium, decreased cytotoxicity **(Fig. 1G, H)**. To test this hypothesis, we used the rhamnolipid biosynthetic mutant MPAO1*rhlA* to measure the lipid content of conditioned medium. This mutant lacked lipid content in its conditioned medium but showed no apparent defects in bacterial growth or pyoverdine production compared to the control strain MPAO1*cat* (transposon inserted in an extraneous gene encoding a chloramphenicol acetyltransferase, **Fig. 2A-C)**, suggesting that much of the lipid material being detected by FM 1-43 was due to rhamnolipid production. Importantly, the conditioned medium from MPAO1*rhlA* neither damaged the 16HBE epithelial monolayer nor induced cell death **(Fig. 2D-G)**. We observed similar results (i.e., reduced lipid content and lower toxicity) for mutants with transposons inserted into genes encoding the RhlRI quorum sensing system (MPAO1*rhlR*, MPAO1*rhlI*) **(Fig. 2A-G)**, indicating that rhamnolipid production is primarily regulated via this signaling pathway in EMEM (30).

**Fig. 2.**
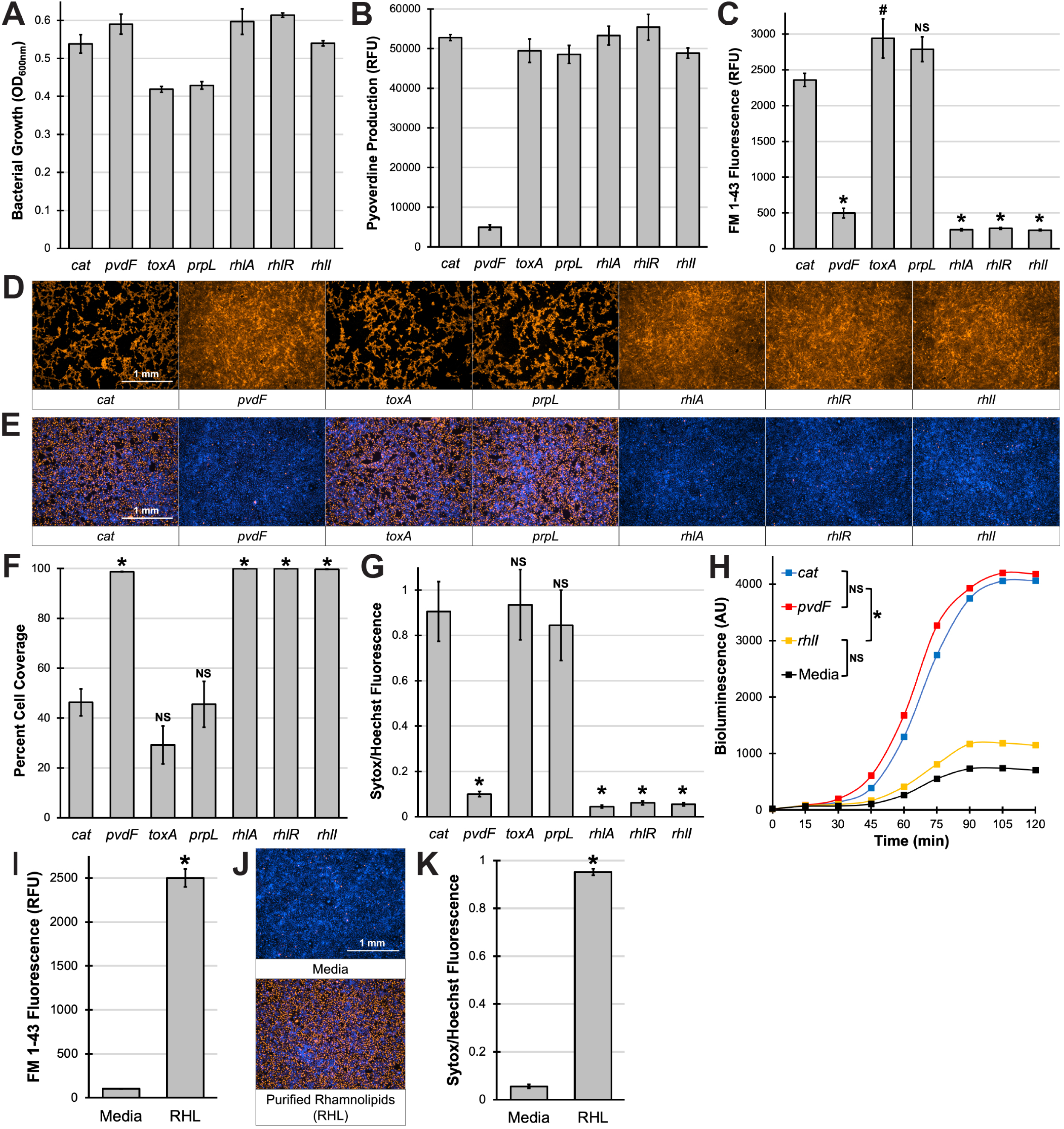
Pyoverdine regulates the production of rhamnolipids. **(A, B)** Bacterial growth **(A)** or pyoverdine production **(B)** by MPAO1 transposon mutants in serum-free EMEM. **(C)** Quantification of lipids in conditioned medium from MPAO1 transposon mutants by FM 1-43 fluorescent labeling. **(D)** Fluorescent micrographs of 16HBE cells after 30 min exposure to conditioned medium from MPAO1 transposon mutants. Cells were prelabeled with CellMask Orange plasma membrane stain. **(E)** Fluorescent micrographs of 16HBE cells after 15 min exposure to conditioned medium from MPAO1 transposon mutants in the presence of Sytox Orange nucleic acid stain (red). Cells were prelabeled with Hoechst 33342 nucleic acid stain (blue). **(F)** Quantification of percentage micrograph area covered by fluorescent cells in **(D)**. **(G)** Quantification of Sytox Orange mean fluorescence intensity normalized to that of Hoechst 33342 in **(E)**. **(H)** Bioluminescence produced by *E. coli* JM109 pSB536 (N-butanoyl-L-homoserine lactone reporter strain) grown in media supplemented with conditioned medium from MPAO1 transposon mutants. **(I)** Quantification of lipids in EMEM with purified rhamnolipids (RHL) by FM 1-43 labeling. **(J)** Fluorescent micrographs of 16HBE cells after 15 min exposure to purified rhamnolipids from *P. aeruginosa* in the presence of Sytox Orange nucleic acid stain (red). Cells were prelabeled with Hoechst 33342 nucleic acid stain (blue). **(K)** Quantification of Sytox Orange mean fluorescence intensity normalized to that of Hoechst 33342 in **(J)**. All error bars represent SEM from three biological replicates. * corresponds to *p* < 0.01, # corresponds to *p* < 0.05, and NS corresponds to *p* > 0.05 based on one-way ANOVA with Dunnett’s multiple comparisons test **(C, F, G)** or Tukey’s multiple comparisons test **(H)** – see **Fig. S4** or Student’s *t*-test **(K)**.

To test whether quorum-sensing is impaired in pyoverdine biosynthetic mutants, we took advantage of an *Escherichia coli*-based bioluminescent reporter that responds to extracellular quorum-sensing molecules (31), specifically N-butanoyl-L-homoserine lactone (C4-HSL) that is produced by the RhlRI system (32). Using this reporter, we quantified C4-HSL concentrations in the conditioned medium of pyoverdine mutants but did not observe significant difference in C4-HSL production between the mutants and their pyoverdine-producing counterparts **(Fig. 2H; Fig. S4)**. For the *rhlI* C4-HSL biosynthetic mutant, the bioluminescent output in the conditioned medium was comparable to that of media control **(Fig. S4)**, indicating that the reporter was selectively responding to extracellular C4-HSL. These results imply that pyoverdine regulates rhamnolipid production through an alternative pathway.

Furthermore, we examined the effects of exposing 16HBE cells to purified rhamnolipids (27). At concentrations comparable to those seen in conditioned medium from wild-type PAO1 or MPAO1*cat* **(Fig. 2I; Fig. S5A)**, purified rhamnolipids were sufficient to kill 16HBE cells **(Fig. 2J, K; Fig. S5C, E)**. However, cells treated with purified rhamnolipids displayed less detachment **(Fig. S5B, D)**, suggesting that other secreted factors (e.g., proteases) in the conditioned medium were required for full damage to the epithelial monolayer. Consistent with these findings, heat denaturation of the conditioned medium did not affect rhamnolipid content **(Fig. S5F)** or cytotoxicity of the material **(Fig. S5H, J),** but it did attenuate cell detachment from the monolayer **(Fig. S5G, I)**. Since the type II secretion system is responsible for the secretion of several *P. aeruginosa* protein toxins and the majority of secreted proteases **(Fig. S3E)** (33), we hypothesized that this system would contribute to cell detachment. We treated 16HBE cells with the conditioned medium from a mutant (MPAO1*xcpQ*) lacking the outer membrane transporter for these secreted proteins but saw no significant attenuation in cell detachment **(Fig. S3A, C)**. It is important to note that while protease production in the type II secretion mutant was substantially impaired, it was not abolished **(Fig. 3E)**. If extracellular proteases were involved in epithelial damage, they could be secreted through other means.

**Fig. 3.**
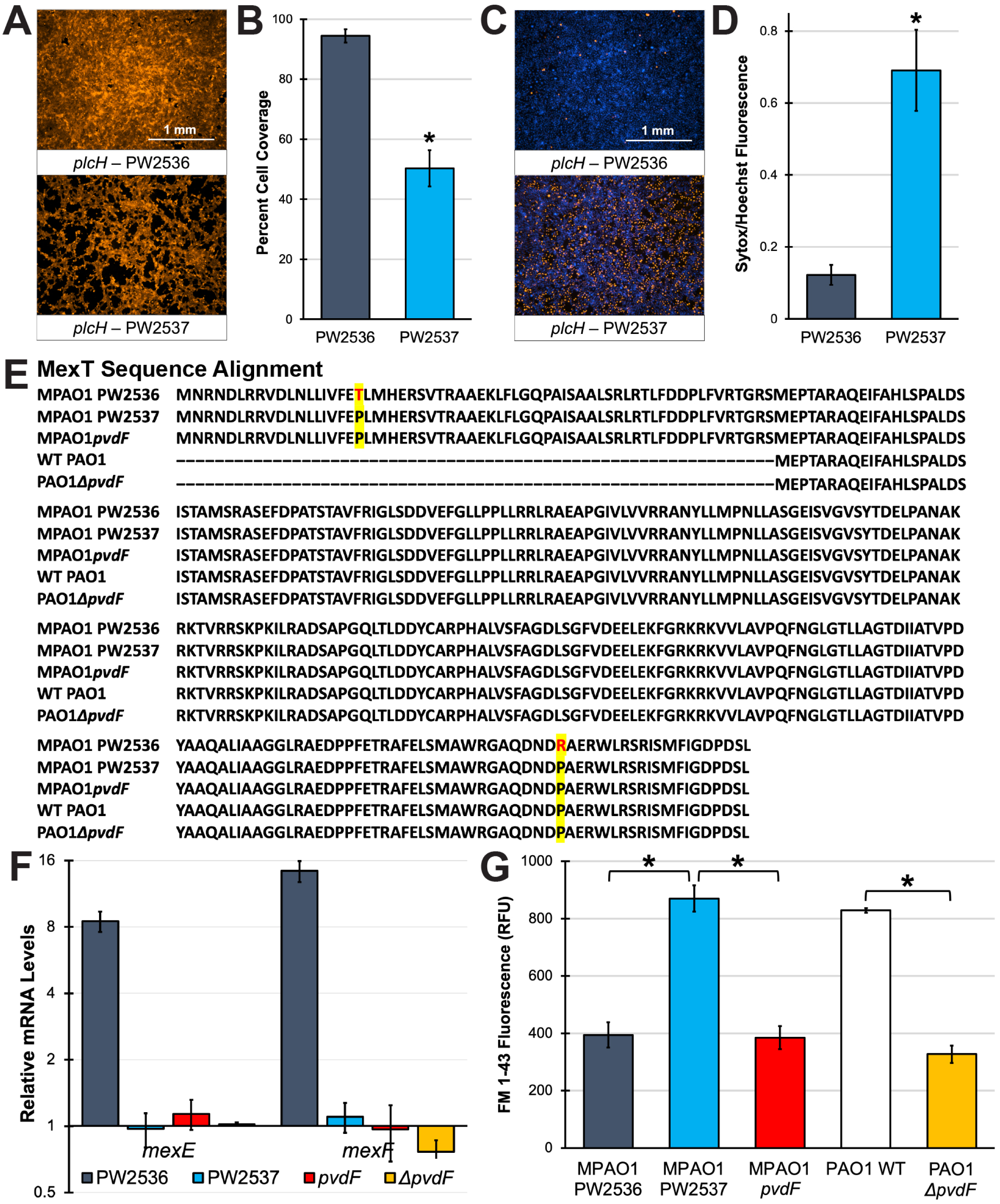
*mexEF* overexpression mutant exhibits decreased rhamnolipid production. **(A)** Fluorescent micrographs of 16HBE cells after 30 min exposure to conditioned medium from *plcH* transposon mutants. Cells were prelabeled with CellMask Orange plasma membrane stain. **(B)** Quantification of percentage micrograph area covered by fluorescent cells. **(C)** Fluorescent micrographs of 16HBE cells after 15 min exposure to conditioned medium from *plcH* transposon mutants in the presence of Sytox Orange nucleic acid stain (red). Cells were prelabeled with Hoechst 33342 nucleic acid stain (blue). **(D)** Quantification of Sytox Orange mean fluorescence intensity normalized to that of Hoechst 33342 in. **(E)** Alignment of MexT protein sequences from MPAO1*plcH* (PW2536), MPAO1*plcH* (PW2537), MPAO1*pvdF*, PAO1, or PAO1*ΔpvdF*. MexT sequences were acquired from paired-end Illumina sequencing (PW2536, PW2537, PAO1, PAO1*ΔpvdF*) or PCR-Sanger sequencing (MPAO1*pvdF*). Amino acid substitution sites are highlighted yellow. Amino acid substitutions in PW2536 are labeled red. **(F)** *mexE* and *mexF* mRNA levels compared to that of WT PAO1. mRNA levels were measured by qRT-PCR. **(G)** Rhamnolipid content in conditioned medium from 14 h EMEM cultures measured by FM 1-43 labeling. All error bars represent SEM from three biological replicates. * corresponds to *p* < 0.01 based on Student’s *t*-test **(B, D)** or one-way ANOVA with Tukey’s multiple comparisons test **(G)**.

### Pyoverdine mutants do not exhibit changes in mexEF expression

While investigating possible secreted toxins that may facilitate rhamnolipid-mediated epithelial damage, we serendipitously identified two presumably isogenic MPAO1 mutants with transposon insertions in *plcH*, a gene that encodes a hemolytic phospholipase C: PW2536 and PW2537. Sanger sequencing verified the presence of the transposon in the *plcH* gene in each of the strains. (PW2536 – 1711/2193nt; PW2537 – 1049/2193nt;). Despite this, incongruous results were obtained when conditioned medium was tested for cytotoxicity against 16HBE cells; conditioned medium from PW2536 showed reduced toxicity, while identically-prepared medium from PW2537 did not **(Fig. 3A-D).** To address this discrepancy, each strain was subjected to whole-genome sequencing, which revealed two monogenic single nucleotide polymorphisms (SNPs) between the two strains. This caused two amino acid substitutions in the transcriptional regulator MexT in PW2536. MexT is known to regulate the *mexEF-oprN* multidrug efflux pump operon **(Fig. 3E)** (34). The mutations in PW2536 led to the overexpression of *mexEF* **(Fig. 3F)**, a phenotype previously associated with decreased rhamnolipid production **(Fig. 3G)** (35, 36).

Based on these observations, we tested whether pyoverdine biosynthetic mutants with poor rhamnolipid production **(Fig. 3G)** had background mutations in MexT or increased *mexEF* transcription. Neither the MPAO1 transposon mutant (MPAO1*pvdF*) nor the PAO1 in-frame deletion mutant (PAO1*ΔpvdF*) had altered MexT protein sequence or changes in *mexEF* expression **(Fig. 3E, F)**, suggesting that pyoverdine regulates rhamnolipid production through an unknown *mexEF*- and C4-HSL- **(Fig. S4)** independent pathway.

### 5-Fluorocytosine inhibits pyoverdine and rhamnolipid production in highly virulent P. aeruginosa clinical isolates

Since genetic disruption of pyoverdine biosynthesis decreased rhamnolipid production, we hypothesized the same could be accomplished using a chemical inhibitor. To that end, we tested whether the pyoverdine biosynthetic inhibitor and FDA-approved antimycotic drug 5-fluorocytosine (5-FC) (23, 37, 38) could inhibit rhamnolipid production in several *P. aeruginosa* strains, including PAO1 and two clinical strains isolated from pediatric CF patients, PA2-72 and PA2-61. These two strains were selected from a large collection of CF isolates for their high *in vitro* pyoverdine production and virulence against the nematode host *C. elegans* **(Fig. 4A, B)** (22). These isolates also exhibited substantial *in vivo* pyoverdine production during acute murine lung infection where they caused host mortality (22). When these strains were grown in EMEM, 5-FC significantly impaired pyoverdine and rhamnolipid production without overtly affecting bacterial growth **(Fig. 4C-F)** with the exception of PA2-61, where the drug induced planktonic cell aggregation **(Fig. 4D)**, confounding bacterial growth measurement by optical density.

**Fig. 4.**
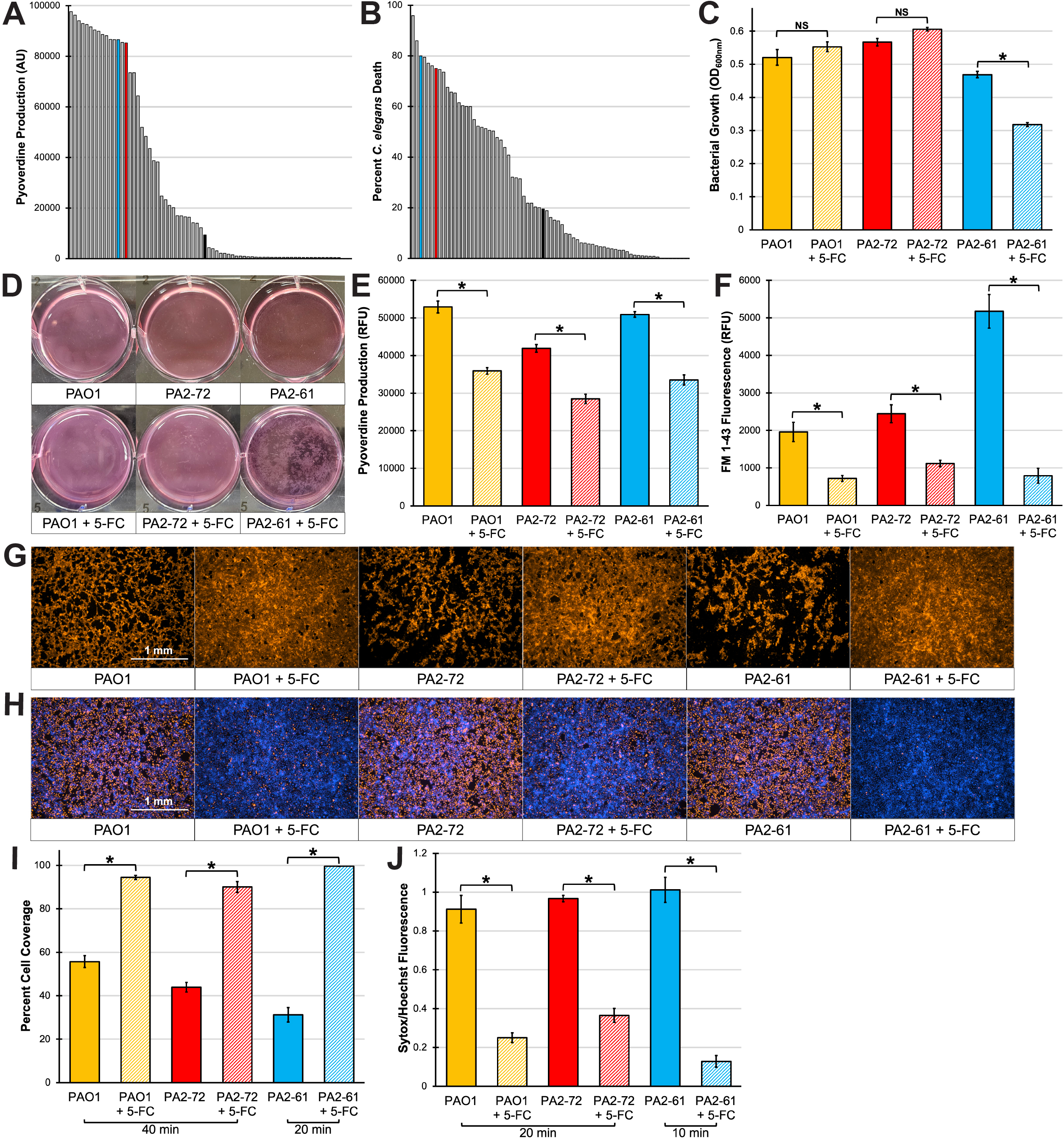
5-Fluorocytosine (5-FC) inhibits pyoverdine and rhamnolipid production in highly virulent cystic fibrosis isolates. **(A, B)** Pyoverdine production **(A)** or *C. elegans* death **(B)** by multidrug-resistant *P. aeruginosa* strains isolated from pediatric cystic fibrosis patients. Black bar represents the median pyoverdine production or *C. elegans death*. Red bar represents PA2-72, blue bar represents PA2-61. Survey data was adapted from (22). **(C)** Bacterial growth of PAO1, PA2-72, or PA2-61 in EMEM with or without 100 µM 5-FC. **(D)** Photograph of EMEM culture after 18 h incubation. **(E, F)** Pyoverdine **(E)** or rhamnolipid **(F)** production of PAO1, PA2-72, or PA2-61 in EMEM with or without 100 µM 5-FC. **(G)** Fluorescent micrographs of 16HBE cells after 40 min (PAO1, PA2-72) or 20 min (PA2-61) exposure to EMEM conditioned medium. Cells were prelabeled with CellMask Orange plasma membrane stain. **(H)** Fluorescent micrographs of 16HBE cells after 20 min (PAO1, PA2-72) or 10 min (PA2-61) exposure to EMEM conditioned medium in the presence of Sytox Orange nucleic acid stain (red). Cells were prelabeled with Hoechst 33342 nucleic acid stain (blue). **(I)** Quantification of percentage micrograph area covered by fluorescent cells in **(G)**. **(J)** Quantification of Sytox Orange mean fluorescence intensity normalized to that of Hoechst 33342 in **(H)**. All error bars represent SEM from four biological replicates. * corresponds to *p* < 0.01 and NS corresponds to *p* > 0.05 based on one-way ANOVA with Sidak’s multiple comparisons test.

5-FC significantly attenuated 16HBE cell detachment and death after exposure to conditioned medium from each of the three strains **(Fig. 4G-J)**. These findings are consistent with previous work where the inhibition of pyoverdine biosynthesis by 5-FC was sufficient to rescue invertebrate and mammalian hosts from *P. aeruginosa* virulence (22, 23, 39).

### Pyoverdine translocates into 16HBE cells but is sequestered within early endosomes

Next, we wanted to investigate the consequences of exposing 16HBE cells to pyoverdine in the absence of other virulence factors. In brief, pyoverdine-rich bacterial filtrate was subjected to two purification steps to separate small molecules by polarity: absorption and elution from a nonpolar polymeric resin (Amberlite XAD-4) and high-performance liquid chromatography (HPLC) via a C-18 reverse-phase column **(Fig. 5A-C)**. We tested whether this purified material was toxic to 16HBE cells using a resazurin-based cell viability assay and compared its toxicity to other known iron-chelating molecules, namely the ferric iron chelator ciclopirox olamine, the ferrous iron chelator 1,10-phenanthroline, or the siderophores pyochelin (from *P. aeruginosa*) or deferoxamine (from *Streptomyces spp.*). While all other iron chelators exhibited time- and dose-dependent cytotoxicity towards 16HBE cells, pyoverdine remained largely nontoxic **(Fig. 5D)** even after 72 h treatment at 200 µM **(Fig. S6A, B)**.

**Fig. 5.**
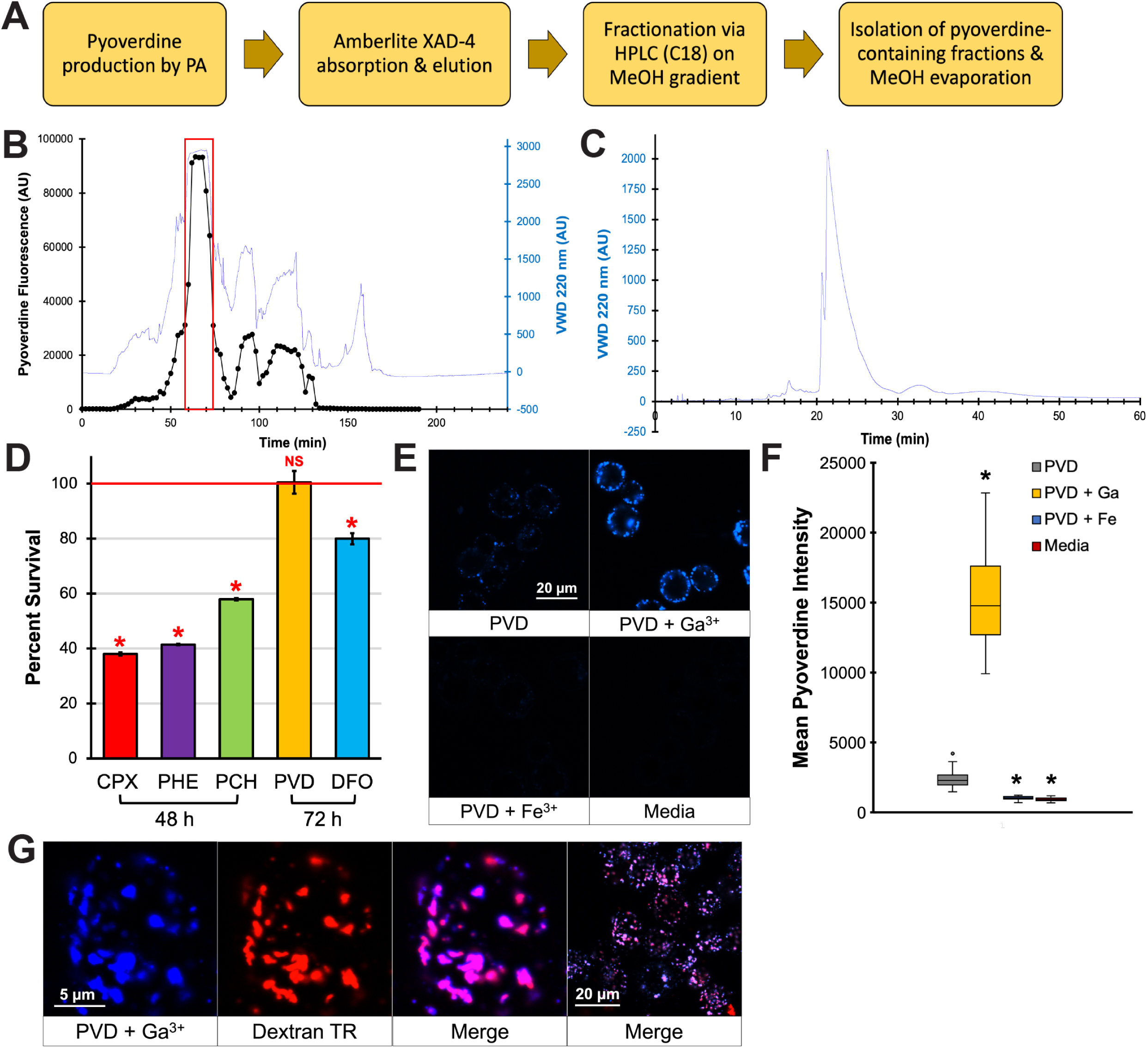
Pyoverdine translocates into 16HBE cells and localizes to early endosomes. **(A)** Summary of the pyoverdine purification pipeline. **(B)** Representative chromatogram from the HPLC purification step of the pipeline. Red box depicts the predominant pyoverdine-containing fractions that were collected. **(C)** Analysis of the final purified product via HPLC. **(D)** 16HBE cell viability after 48 h treatment with 100 µM ciclopirox olamine, 1, 10-phenanthroline, or pyochelin, or 72 h treatment of 100 µM pyoverdine or deferoxamine in serum-free EMEM. **(E)** Confocal micrographs of 16HBE cells exposed to 100 µM purified pyoverdine, pyoverdine with excess Ga(NO_3_)_3,_ pyoverdine with excess FeCl_3_, or media control for 24 h. Cells were trypsinated prior to imaging. **(F)** Quantification of pyoverdine fluorescence within 30 individual cells. **(G)** Confocal micrographs of 16HBE cells treated with 100 µM pyoverdine-gallium and dextran-Texas Red (10,000 MW). Error bars in **(D)** represent SEM from three biological replicates. Error bars in **(C)** represent standard deviation. * corresponds to *p* < 0.01 based on one-way ANOVA with Dunnett’s multiple comparisons test.

Because pyoverdine is considerably larger than these iron chelators, with a molecular weight of ∼1,365 g/mol, it may be unable to translocate across cellular membranes. We took advantage of pyoverdine’s intrinsic spectral properties to examine whether pyoverdine could translocate into 16HBE cells. After 24 h, 16HBE cells treated with purified pyoverdine showed considerable internalization of pyoverdine **(Fig. 5E, F)**. Consistent with previous studies, this intracellular fluorescence was enhanced when pyoverdine was presaturated with gallium and was quenched when pyoverdine was presaturated with iron **(Fig. 5E, F)** (21, 26). Importantly, pyoverdine fluorescence did not colocalize with that of the plasma membrane stain, indicating that pyoverdine was within the cell **(Fig. S6C)**. However, we also observed that pyoverdine fluorescence formed distinct punctae within the cell. Based on previous observations in murine macrophages (26), we hypothesized that pyoverdine was sequestered within early endosomes. Supporting this hypothesis, pyoverdine colocalized with fluorophore-conjugated 10 kDa dextran, a well-established endosomal marker **(Fig. 5G, Fig. S6D)** (40). These results are consistent with our observations that pyoverdine, unlike other iron chelating molecules, exhibited low cytotoxicity towards 16HBE cells.

### Iron chelation activates a proinflammatory response in 16HBE cells

While pyochelin exhibits lower affinity towards ferric iron than pyoverdine, it is also substantially smaller, with a molecular weight of ∼325 g/mol. We hypothesized that pyochelin may be able to enter 16HBE cells and chelate intracellular iron. While we were not able to visualize pyochelin within cells (due to its lack of distinct spectral properties), one likely consequence of iron deprivation in epithelial cells would be a proinflammatory transcriptional response. Several studies have demonstrated that iron chelation by various siderophores such as deferoxamine or enterobactin promotes the production of proinflammatory cytokines, most notably interleukin (IL)-8 in lung epithelial cells, intestinal epithelial cells, or oral keratinocytes (41–43). To reaffirm these findings, we treated 16HBE cells with various iron chelators and measured the mRNA levels of genes involved in neutrophilic inflammation. We first observed that total RNA yield (from phenol-chloroform extraction) in these cells corresponded with the resazurin-based cell viability assay **(Fig. 5D)**. Cells treated with small molecule (< 1,000 MW), cytotoxic iron chelators (ciclopirox olamine, phenanthroline, pyochelin, or deferoxamine) yielded lower quantities of RNA, while cells treated with pyoverdine had RNA quantities comparable to that of media control **(Fig. 6A)**. Using qRT-PCR, we measured the expression of genes encoding components of the inflammasome, namely *NLRP3* and *NLRP1*, and those encoding the major proinflammatory cytokines produced by lung epithelial cells, specifically IL-1β (*IL1B*), IL-8 (*IL8*), and tumor necrosis factor alpha (*TNF*). All of these genes have been associated with inflammation during lung infection (44). With the exception of pyoverdine, all iron chelators induced the expression of these proinflammatory genes **(Fig. 6B)**. For *IL8*, we validated the qRT-PCR results by ELISA to determine whether transcriptional activation led to increased cytokine production. Cells treated with cytotoxic iron chelators exhibited time-dependent increase in IL-8 secretion that correlated to *IL8* mRNA levels **(Fig. 6C, Fig. S7)**. Cells treated with pyoverdine exhibited IL-8 secretion comparable to that of media control **(Fig. 6C)**. To ensure that the observed proinflammatory response was due to iron chelation, we presaturated pyochelin and deferoxamine with excess gallium (1:2 stoichiometric ratio) prior to exposure. Cells treated with gallium-bound pyochelin or deferoxamine yielded RNA quantities comparable to that of media control, suggesting that gallium inhibited the cytotoxic effects of the siderophores **(Fig. 6D)**. Furthermore, pretreating the siderophores with gallium resulted in a significant decrease in proinflammatory gene expression **(Fig. 6E)**, demonstrating that the siderophore-induced inflammatory response was due to iron chelation rather than other nonspecific reactions or contaminants in the commercially-sourced material.

**Fig. 6.**
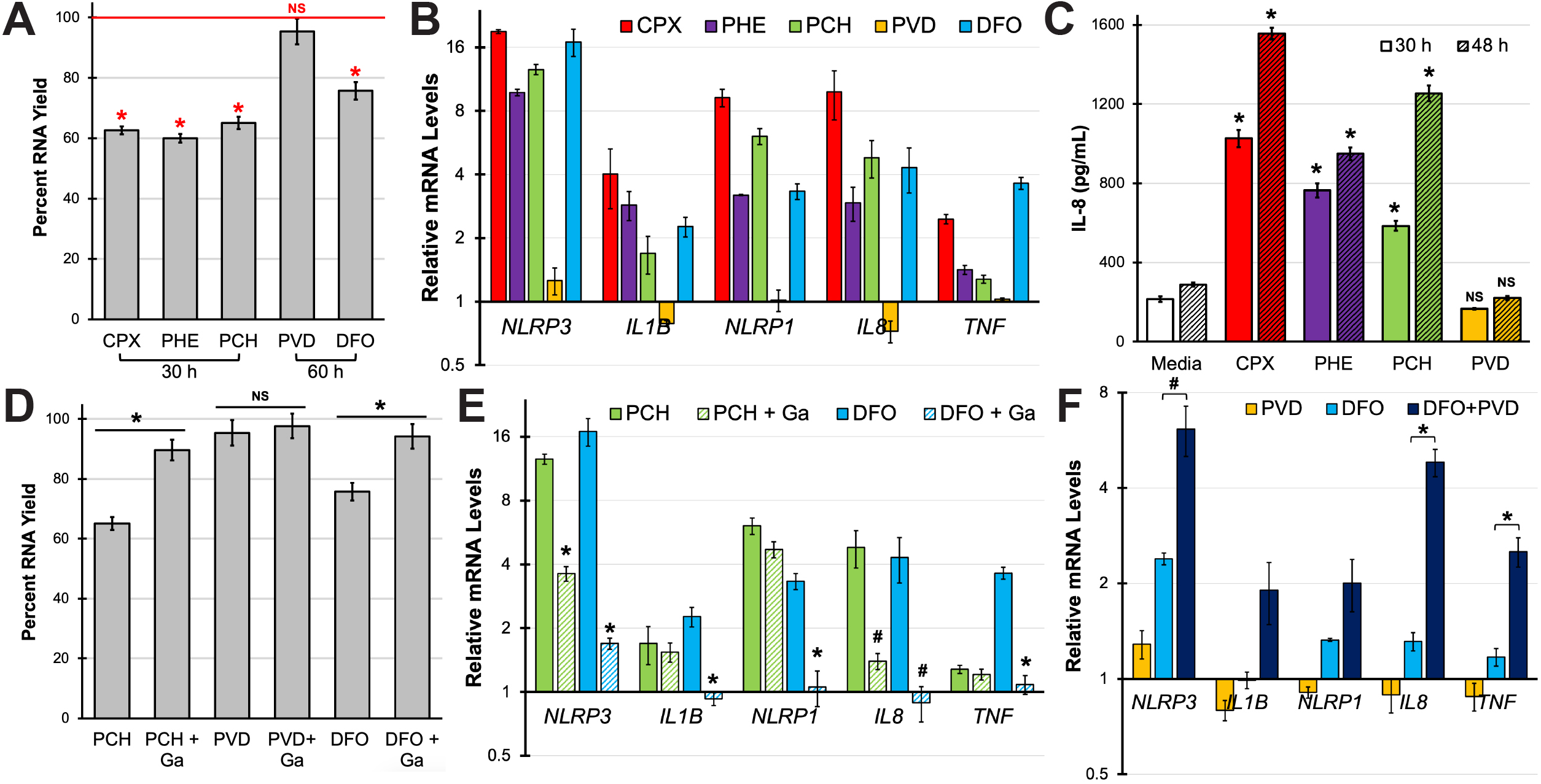
Small molecule iron chelators promote the expression of proinflammatory genes in 16HBE cells. **(A)** Total RNA yield in 16HBE cells treated with ciclopirox olamine (CPX), 1,10-phenanthroline (PHE), or pyochelin (PCH) for 30 h or cells treated with pyoverdine (PVD) or deferoxamine (DFO) for 60 h normalized to that of media control. All treatments were at 100 µM in serum-free EMEM. **(B)** Proinflammatory gene expression (*NLRP3, IL1B, NLRP1, IL8, TNF*) in cells treated with iron chelators compared to that of media control. mRNA levels were measured by qRT-PCR. **(C)** IL-8 protein concentration in the supernatants of 16HBE cells treated with iron chelators. IL-8 was quantified by ELISA. **(D, E)** Total RNA yield **(D)** or proinflammatory gene expression **(E)** in 16HBE cells treated with iron chelators with or without excess Ga(NO_3_)_3_ supplementation. **(F)** Proinflammatory gene expression after 60 h treatment with pyoverdine, deferoxamine, or both molecules. All error bars represent SEM from three biological replicates. * corresponds to *p* < 0.01, # corresponds to *p* < 0.05, and NS corresponds to *p* > 0.05 based on one-way ANOVA with Dunnett’s **(A, C)**, Sidak’s **(D, E)**, or Tukey’s **(F)** multiple comparisons test.

Finally, we investigated whether pyoverdine can indirectly promote lung inflammation by potentiating other iron chelating molecules. Due to its exceptionally high affinity for iron, pyoverdine is likely to remove iron from other, more cell-permeable siderophores or outcompete them for trace iron in the extracellular milieu, increasing the pool of apo-siderophores that can promote inflammation. To test this hypothesis, we treated 16HBE cells with deferoxamine, pyoverdine, or both. Cells treated with both siderophores exhibited higher expression of proinflammatory genes compared to those treated with deferoxamine alone **(Fig. 6F)**. Considering that pyoverdine alone did not affect the transcription of proinflammatory genes, these results suggest that pyoverdine enhanced deferoxamine-mediated damage by removing iron from deferoxamine and effectively increasing the pool of apo-deferoxamine.

## Discussion

Arguably one of the greatest challenges to combating *P. aeruginosa* infections is the sheer multitude of virulence factors produced by the bacterium that contribute to pathogenesis. These include small molecule virulence factors (e.g., siderophores, quorum-sensing molecules), factors involved in biofilm formation and motility (e.g, exopolysaccharides, type IV pili, flagella), and more than twenty toxins that either directly kill host cells (e.g., exotoxin A, exoenzyme S, exotoxin T, exotoxin U) or damage host tissue (e.g., elastase LasA, elastase LasB, PrpL, alkaline protease) (32, 45–48). This complexity casts a dark shadow over the prospects of epidemiological or therapeutic intervention. Ideally, we would be able to reliably predict a pathogen’s ability to cause disease through our evolving molecular surveillance tools such as whole-genome sequencing and mass spectrometry and therapeutically impair pathogenesis through antivirulence drugs that inhibit the production or function of key virulence factors and toxins. In *P. aeruginosa*, the only feasible way to approach these strategies would be to unravel the regulation of virulence factors in the bacterium and to target virulence networks rather than individual factors.

The results we report in this study suggest that the alternative sigma factor PvdS may be a promising target for therapeutic intervention during *P. aeruginosa* lung infections. PvdS has already been shown to regulate the production of several secreted toxins, such as the translational inhibitor exotoxin A and the secreted protease PrpL. Exotoxin A, arguably one of the most extensively studied toxins in *P. aeruginosa*, inhibits protein synthesis (46, 49), inducing airway epithelial cell death (50) and inhibiting cell junction repair in the presence of *P. aeruginosa* elastase (51). Exotoxin A also contributes to *P. aeruginosa* virulence in various murine infection models (52–54). PrpL degrades host defense factors, like surfactant proteins and IL-22, that contribute to lung innate immunity (55–57). PrpL has also been shown to directly contribute to *P. aeruginosa* virulence during ocular infections (58, 59).

PvdS is best known for its role in pyoverdine biosynthesis and is indispensable for the production of pyoverdine biosynthetic enzymes. In addition to scavenging trace iron in the environment or directly from host ferroproteins, pyoverdine is involved in a positive feedback loop where the uptake of iron-bound pyoverdine by its outer membrane receptor, FpvR, derepresses PvdS, increasing the production of pyoverdine, exotoxin A, and PrpL (12, 18). Here, we have demonstrated that pyoverdine promotes the production of an additional secreted material, rhamnolipids, that rapidly induce cell death. Secreted rhamnolipids have been shown to assemble into micellar structures (60) that directly interact with host membranes, causing rapid plasma membrane rupture and cell death (29, 61, 62) and damaging the lung epithelium (63, 64). It remains unclear however how pyoverdine regulates rhamnolipid production and whether this mechanism is linked to quorum-sensing, the primary mode of rhamnolipid regulation in the bacterium. While certain studies suggest that *P. aeruginosa* quorum-sensing affects pyoverdine production (65–69), an inverse relationship has yet to be explored.

In addition to regulating secreted toxins, pyoverdine may also indirectly contribute to inflammation by removing iron from other more cell permeable siderophores such as pyochelin, deferoxamine, or enterobactin, the latter in the context of polymicrobial infections with *Enterobacteriaceae* such as the respiratory pathogen *K. pneumoniae* (70). Importantly, while *P. aeruginosa* may lose the ability to produce pyoverdine during lung infection with the emergence of social cheaters or due to a transition in iron acquisition strategy (71–73), several surveys of patient sputum samples and clinical isolates have revealed that a large fraction of strains still exhibit substantial pyoverdine production (22, 74–76), demonstrating that pyoverdine may be an important target for therapeutic intervention.

We took advantage of the FDA-approved antimycotic drug, 5-fluorocytosine (5-FC), that inhibits *pvdS* expression in *P. aeruginosa* and attenuates virulence during murine lung infection. Imperi and colleagues first identified 5-FC in a screen for small molecules that inhibit pyoverdine production (23). We have independently identified a chemical analogue of 5-FC, 5-fluorouracil – another *pvdS* inhibitor, in a small molecule screen for compounds that rescue *C. elegans* from *P. aeruginosa* in a pyoverdine-dependent pathogenesis model (37, 77). We also recently reported that 5-FC synergizes with another FDA-approved drug, gallium nitrate, to inhibit *P. aerugin*osa growth and virulence against *C. elegans* (38). Our findings in this study suggest that in addition to its bactericidal and biofilm-inhibitory activities (78, 79), gallium could also function as an anti-inflammatory agent during lung infection by inhibiting intracellular iron chelation by pyochelin and mitigating not only epithelial cell death, but also activation of proinflammatory pathways such as the NLRP3 inflammasome and IL-8 production. These newly discovered roles for pyochelin further demonstrate that pyoverdine and pyochelin play distinct roles in *P. aeruginosa* virulence. Previous studies have shown that pyochelin triggers reactive oxygen species (ROS) production in an iron-dependent manner in host cells during infection (80) or in other microbes during interbacterial competition (81–83). While pyochelin may not regulate additional virulence pathways, its ability to permeate membranes makes this siderophore a distinctly effective tool for direct host cell damage.

The benefits of suppressing pyochelin-mediated neutrophilic inflammation during lung infection, particularly chronic lung infection, has been well documented. While the mechanisms neutrophils employ to kill and remove pathogens such as the production of neutrophil elastases are important for host defense, they can also cause tissue damage by degrading extracellular matrix proteins (84, 85). During chronic infections (such as those in CF patients), these host defense factors continue to cause airway damage while the pathogen persists, exacerbating the decline in pulmonary function (86). While lung inflammation is mediated by many factors (cytokines and chemokines), for CF patients, a strategy to specifically inhibit the NLRP3 inflammasome by therapeutics such as MCC950 is currently being investigated with promising results in murine infection studies (87, 88). It is important to note that NLRP3 inflammasome priming (i.e., transcriptional upregulation of *NLRP3*, encoding the major component of the inflammasome, and *IL1B*, encoding pro-IL-1β) (89) was a pathway that was activated by intracellular iron chelation in lung epithelial cells **(Fig. 6B)**. We observed this transcriptional response in not only wild-type 16HBE cells but also those carrying mutations in the cystic fibrosis transmembrane conductance regulator (CFTR G551D, CFTR ΔF508), two of the most frequently identified mutations in CF patients **(Fig. S8)** (90). While gallium has been broadly associated with anti-inflammatory properties (91–93), studies have yet to specifically explore gallium’s role in inhibiting pathogen-associated inflammation. Considering recent findings that bacterial siderophores promote inflammation (41–43, 70), this therapeutic avenue may merit consideration.

## Materials and Methods

### Bacterial Strains and Growth Conditions

See list of bacterial strains in **Table 1**. All MPAO1 transposon insertion sites were verified by Sanger sequencing (94). Insertions were determined by adapting a previously established method for the MAR2xT7 transposon library in PA14 (95) and primers in **Table S2**. To produce pyoverdine-rich conditioned medium, an LB overnight culture of *P. aeruginosa* was diluted 20-fold into 2 mL of serum-free Eagle’s Minimum Essential Medium (EMEM) (Millipore Sigma, St. Louis, MO) in a 6-well plate. The plate was sealed with a Breathe-Easy sealing membrane (Diversified Biotech, Dedham, MA) and grown statically at 37 ℃ for 18 h. Pyoverdine production (Ex. 405 nm; Em. 460 nm) and bacterial growth (Abs. 600 nm) were measured spectrophotometrically on a Cytation5 Multimode Reader (Biotek, Winnoski, VT). Bacteria was then removed by centrifugation and the supernatant was treated with an antibiotic cocktail to kill residual bacteria (100 µg/mL amikacin, 100 µg/mL gentamicin, 100 µg/mL tobramycin).

**Table 1.** List of bacterial strains and cell lines used in this study.

| Strains | Relevant Genotype | Source or Reference |
| --- | --- | --- |
| *Pseudomonas aeruginosa* Strains |  |  |
| PAO1 | WT | D. Haas |
| PAO1ΔpvdF | ΔpvdF | D. Haas |
| PAO1ΔpchBA | ΔpchBA | D. Haas |
| PAO1ΔpvdFpchBA | ΔpvdFpchBA | D. Haas |
| MPAO1 PW2280 | cat (ISlacZ/hah) | Jacobs et al., 2003 |
| MPAO1 PW5033 | pvdF (ISlacZ/hah) | Jacobs et al., 2003 |
| MPAO1 PW3078 | toxA (ISphoA/hah) | Jacobs et al., 2003 |
| MPAO1 PW8077 | prpL (ISphoA/hah) | Jacobs et al., 2003 |
| MPAO1 PW6887 | rhlA (ISphoA/hah) | Jacobs et al., 2003 |
| MPAO1 PW6882 | rhlR (ISlacZ/hah) | Jacobs et al., 2003 |
| MPAO1 PW6880 | rhlI (ISphoA/hah) | Jacobs et al., 2003 |
| MPAO1 PW5085 | pvdS (ISlacZ/hah) | Jacobs et al., 2003 |
| MPAO1 PW6223 | xcpQ (ISphoA/hah) | Jacobs et al., 2003 |
| MPAO1 PW2537 | plcH (ISlacZ/hah) | Jacobs et al., 2003 |
| MPAO1 PW2536 | plcH (ISlacZ/hah) | Jacobs et al., 2003 |
| PA2-72 | WT (CF Isolate) | Kang et al., 2019 |
| PA2-61 | WT (CF Isolate) | Kang et al., 2019 |
| *Other Bacterial Strains* |  |  |
| *Escherichia coli* JM109 pSB536 | *PahyI::luxABCDE* regulated by constitutive AhyR | Winson et al., 1998 |
| *Human Bronchial Epithelial Cells* |  |  |
| 16HBE | WT CFTR | G. Bao |
| 16HBE G551D | CFTR G551D | G. Bao |
| 16HBE ΔF508 | CFTR ΔF508 | G. Bao |

### Cell Culture

Wild-type and mutant 16HBE cells **(Table 1)** were passaged in Eagle’s Minimum Essential Medium (EMEM) supplemented with 10% fetal bovine serum (Corning, Corning, NY), penicillin/streptomycin (Millipore Sigma), and MEM non-essential amino acids (Millipore Sigma). For experiments with *P. aeruginosa* conditioned medium, 4⨉10^6^ cells were seeded into each well of a collagen (type I from calf skin - Millipore Sigma)-coated 12-well plate and grown at 37 ℃ for 24 h in a CO_2_-jacketed incubator by which they reached 100% confluence. To visualize the epithelial monolayer, cells were stained with 2.5 µg/mL CellMask Orange plasma membrane stain (Invitrogen, Carlsbad, CA) for 1 h prior to conditioned medium exposure. Following treatment, the medium was aspirated and the monolayer was imaged on a Cytation5 Multimode Reader using a RFP filter cube. Percentage image area covered by fluorescent cells was quantified using ImageJ. To visualize cell death, cells were prelabeled with 20 µM Hoechst 33342 (ThermoFisher Scientific, Waltham, MA) for 30 min then exposed to conditioned medium in the presence of 2.5 µM Sytox Orange (Invitrogen). Following treatment, the medium was aspirated, and the monolayer was imaged on a Cytation5 Multimode Reader using DAPI (for Hoechst 33342) and RFP (for Sytox Orange) filter cubes. Images were exported and quantified for mean blue or red fluorescence intensity on ZEN Blue image analysis software (Zeiss, Oberkochen, Germany).

For 16HBE cell viability measurements following iron chelator treatment, 440 µM resazurin (ThermoFisher Scientific) in phosphate buffered saline was diluted 10-fold into the treatment medium, and cells were incubated for 1.5 h. The medium was collected and briefly centrifuged to remove cells. 150 µL of the supernatant was transferred to a 96-well plate, and resorufin (reduced resazurin) fluorescence (Ex. 560 nm; Em. 590 nm) was measured on a Cytation5 Multimode Reader.

### N-butanoyl-L-homoserine lactone (C4-HSL) Measurement

To measure C4-HSL concentration, one part spent culture medium from *P. aeruginosa* grown in EMEM was mixed with three parts *E. coli* JM109 pSB536 in LB medium. This LB medium was inoculated with the *E. coli* reporter strain by diluting an overnight culture 5-fold. 150 µL of the *P. aeruginosa* conditioned medium-reporter strain mixture was transferred to each well in a 96-well plate and incubated at 37 ℃ for 2 h in a Cytation5 Multimode Reader. Bioluminescence measurements were taken every 15 min.

### P. aeruginosa whole genome sequence analysis

Bacterial genomic DNA was purified from overnight culture using DNeasy UltraClean Microbial Kit (Qiagen, Hilden, Germany). Paired end Illumina sequencing was performed by the Microbial Genome Sequencing Center (MiGS, Pittsburgh, PA) for at least 40X genome coverage. To compare the two *plcH* transposon mutants, raw sequencing reads from PW2537 were first assembled via SPAdes (96) and annotated via Prokka using PAO1 (NC_002516.2) as the reference genome (97). Genomic polymorphisms in PW2536 were identified using *breseq* (98).

### Pyoverdine Purification

A LB overnight culture of *P. aeruginosa* PAO1 was diluted 100-fold into 300 mL of M9 medium (1% w/v 5X M9 Salts (BD Difco, Franklin Lakes, NJ), 1.5% w/v Bacto Casamino Acids with low iron and salt content (BD Difco), 1 mM MgSO_4_, 1 mM CaCl_2_) in a 2 L flask and grown aerobically for 24 h at 37 ℃. Bacteria were then removed by centrifugation and filtration through a 0.22 µm membrane. The filtrate was incubated with 10% w/v amberlite XAD-4 resin (MilliporeSigma) at room temperature for 4 h with constant agitation. After rinsing the resin with copious amounts of water, pyoverdine was eluted in 50% methanol. This eluent was diluted in water to 15% methanol and loaded onto a Luna Omega 5 µm Polar C18 LC prep column (Phenomenex, Torrance, CA) for high-performance liquid chromatography on a 1220 Infinity LC system (Agilent Technologies, Santa Clara, CA). Pyoverdine was eluted from the column by a 0-100% methanol gradient across 4 h at a flowrate of 5 mL/min. Fractions were collected every other minute for pyoverdine content analysis **(Fig. 5B)**. The fractions with the highest pyoverdine content were pooled. Methanol was evaporated using a SpeedVac vacuum concentrator. The final purified product was analyzed by HPLC on an analytical column to verify sample purity **(Fig. 5C)**.

### Confocal Laser Scanning Microscopy

8⨉10^6^ 16HBE cells were seeded into each well of a collagen-coated 6-well plate and grown at 37 ℃ for 24 h in a CO_2_-jacketed incubator by which they reached 100% confluence. After treatment, cells were washed in serum-free EMEM and detached from the microtiter plate by trypsin-EDTA solution (MilliporeSigma). After inactivating the trypsin with media containing 10% fetal bovine serum, the cells were concentrated via centrifugation and transferred onto a glass side with a 3% noble agar pad. These slides were visualized under a LSM800 AiryScan confocal laser scanning microscope (Zeiss). Pyoverdine fluorescence was visualized via a 405 nm laser line using the channel conditions for Pacific Blue. Dextran-Texas Red (Invitrogen) fluorescence was visualized via a 561 nm laser line using channel conditions for Texas Red. CellMask Deep Red plasma membrane stain (Invitrogen) fluorescence was visualized via a 640 nm laser line using channel conditions for Alex Fluor 660.

### qRT-PCR

For *P. aeruginosa*, bacterial cells were collected from 12 mL of EMEM culture by centrifugation. The pellet was resuspended in 2 mL of TRI reagent (Molecular Research Center, Cincinnati, OH) for phenol/chloroform/guanidinium thiocyanate RNA extraction according to manufacturer’s protocols (bromochloropropane phase separation followed by isopropanol RNA precipitation). Prior to phase separation, bacterial cells were homogenized with 0.1 mm zirconia beads by vigorous vortexing. To remove DNA contaminants in bacterial RNA extracts, samples were treated with DNase I (ThermoFisher Scientific) at 37 ℃ for 30 min, followed by 75 ℃ enzyme heat denaturation for 10 min. For 16HBE cells grown and treated in 6-well plates, the treatment medium was aspirated, and cells were incubated in TRI reagent at room temperature for 15 min to lyse cells prior to RNA extraction.

For both bacterial and human cell RNA, cDNA synthesis was performed on a Bio-Rad T100 Thermo Cycler (Bio-Rad, Hercules, CA) using a reverse transcription kit (Applied Biosystems, Waltham, MA). qRT-PCR was performed on a Bio-Rad CFX Connect Real-Time System (Bio-Rad) using a universal qPCR master mix (New England Biolabs, Ipswich, MA). All qPCR primer sequences are in **Table S2**. For *P. aeruginosa* genes, cDNA amplification (Ct value) was normalized to that of housekeeping gene *gyrB*. For 16HBE genes, cDNA amplification was normalized to that of *ACTB*.

## Acknowledgements

This study was supported by funding from the Cystic Fibrosis Foundation (KIRIEN20I0 to NVK; XU23H0 to QX; KANG19H0, KANG22H0 to DK), National Institutes of Health (R35GM129294 to NVK), and American Heart Association (903591 to DK). The authors declare no conflict of interest.

## Supplemental Figure Legends

**Fig. S1. Disruption of pyochelin biosynthesis does not mitigate epithelial damage. (A, B)** Bacterial growth **(A)** or pyoverdine production **(B)** of *P. aeruginosa* PAO1 siderophore biosynthetic mutants (PAO1*ΔpvdF* – pyoverdine; PAO1*ΔpchBA* – pyochelin; PAO1*ΔpvdFΔpchBA* – pyoverdine and pyochelin) after 18 h growth in serum-free EMEM. **(C)** MH-S murine alveolar macrophage viability after 2 h exposure to conditioned medium from PAO1 siderophore mutants grown in EMEM. Cell viability was measured using a resazurin-based assay. Fluorescent micrographs of 16HBE cells after 30 min exposure to conditioned medium from PAO1 siderophore mutants. Cells were prelabeled with CellMask Orange plasma membrane stain. Quantification of percentage micrograph area covered by fluorescent cells. All error bars represent SEM from four biological replicates. * corresponds to *p* < 0.01 and NS corresponds to *p* > 0.05 based on one-way ANOVA with Tukey’s multiple comparisons test.

**Fig. S2. Lipid extraction abrogates epithelial damage by pyoverdine-rich conditioned medium. (A)** Fluorescent micrographs of 16HBE cells after 30 min exposure to conditioned medium from WT PAO1. Conditioned medium was pretreated with 200 µM Ga(NO_3_)_3_, 200 µM FeCl_3_, or 100 µg/mL proteinase K for 24 h, had macromolecules depleted via a 10 kDa centrifugal filter, or had lipids depleted by chloroform (CHCl_3_) extraction. Cells were prelabeled with CellMask Orange plasma membrane stain. **(B)** Quantification of percentage micrograph area covered by fluorescent cells. Error bars represent SEM from three biological replicates. * corresponds to *p* < 0.01 and NS corresponds to *p* > 0.05 based on one-way ANOVA with Dunnett’s multiple comparisons test.

**Fig. S3. Type II secretion system toxins do not contribute to lung epithelial damage. (A)** Fluorescent micrographs of 16HBE cells after 30 min exposure to conditioned medium from MPAO1 transposon mutants. Cells were prelabeled with CellMask Orange plasma membrane stain. Fluorescent micrographs of 16HBE cells after 15 min exposure to conditioned medium from MPAO1 transposon mutants in the presence of Sytox Orange nucleic acid stain (red). Cells were prelabeled with Hoechst 33342 nucleic acid stain (blue). **(C)** Quantification of percentage micrograph area covered by fluorescent cells in **(A)**. **(D)** Quantification of Sytox Orange mean fluorescence intensity normalized to that of Hoechst 33342 in **(B)**. **(E)** Protease activity in conditioned medium from MPAO1 transposon mutants. Proteolytic activity was measured by fluorescence release from cleavage of FITC-conjugated casein. All error bars represent SEM from three biological replicates. * corresponds to *p* < 0.01 and NS corresponds to *p* > 0.05 based on one-way ANOVA with Dunnett’s multiple comparisons test.

**Fig. S4. Pyoverdine biosynthetic mutants do not exhibit impaired C4-HSL quorum-sensing. (A)** Bioluminescence produced by *E. coli* JM109 pSB536 (N-butanoyl-L-homoserine lactone reporter strain) grown in media supplemented with conditioned medium from wild-type PAO1 or PAO1*ΔpvdF*. **(B)** Bioluminescence at the kinetics **(A)** endpoint. **(C)** Bioluminescence produced by the *E. coli* reporter strain grown in media supplemented with conditioned medium from MPAO1 transposon mutants. **(D)** Bioluminescence at the kinetics **(C)** endpoint. All error bars represent SEM from at least three biological replicates. * corresponds to *p* < 0.01 and NS corresponds to *p* > 0.05 based on one-way ANOVA with Tukey’s multiple comparisons test.

**Fig. S5. Purified rhamnolipids are sufficient for 16HBE cell death but not for epithelial monolayer damage. (A)** Quantification of rhamnolipids in purified samples (RHL) or *P. aeruginosa* conditioned medium from WT PAO1, PAO1*ΔpvdF*, MPAO1*cat*, or MPAO1*pvdF.* **(B)** Fluorescent micrographs of 16HBE cells after 30 min exposure to purified rhamnolipids in EMEM. Cells were prelabeled with CellMask Orange plasma membrane stain. **(C)** Fluorescent micrographs of 16HBE cells after 15 min exposure to purified rhamnolipids in EMEM in the presence of Sytox Orange nucleic acid stain (red). Cells were prelabeled with Hoechst 33342 nucleic acid stain (blue). **(D)** Quantification of percentage micrograph area covered by fluorescent cells in **(B)**. **(E)** Quantification of Sytox Orange mean fluorescence intensity normalized to that of Hoechst 33342 in **(C)**. **(F)** Quantification of rhamnolipids in conditioned medium from WT PAO1 or conditioned medium after heat denaturation (85 ℃ for 1 h). **(G)** Fluorescent micrographs of 16HBE cells after 30 min exposure to conditioned medium from WT PAO1 or conditioned medium after heat denaturation. Cells were prelabeled with CellMask Orange plasma membrane stain. **(H)** Fluorescent micrographs of 16HBE cells after 15 min exposure to conditioned medium from WT PAO1 or conditioned medium after heat denaturation in the presence of Sytox Orange nucleic acid stain (red). Cells were prelabeled with Hoechst 33342 nucleic acid stain (blue). **(I)** Quantification of percentage micrograph area covered by fluorescent cells in **(G)**. **(J)** Quantification of Sytox Orange mean fluorescence intensity normalized to that of Hoechst 33342 in **(H)**. All error bars represent SEM from three biological replicates. # corresponds to *p* < 0.05 and NS corresponds to *p* > 0.05 based on Student’s *t*-test.

**Fig. S6. Pyoverdine accumulates in early endosomes of lung epithelial cells. (A, B)** 16HBE cell viability after 48 **(A)** or 72 h **(B)** treatment with ciclopirox olamine, 1, 10-phenanthroline, pyoverdine, or deferoxamine in serum-free EMEM. **(C)** Confocal micrographs of 16HBE cells exposed to 100 µM purified pyoverdine, pyoverdine with excess Ga(NO_3_)_3,_ pyoverdine with excess FeCl_3_, or media control for 24 h. Cells were labeled with CellMask Deep Red plasma membrane stain and trypsinated prior to imaging. **(D)** Confocal micrographs of 16HBE cells treated with 100 µM pyoverdine-gallium and dextran-Texas Red (10,000 MW). Bottom row shows an enlarged micrograph of one representative cell. All error bars represent SEM from three biological replicates.

**Fig. S7. Small molecule iron chelators promote IL-8 production in 16HBE cells. (A)** *IL8* mRNA levels in 16HBE cells treated with iron chelators (ciclopirox olamine – CPX, 1,10-phenanthroline – PHE, pyochelin – PCH, pyoverdine – PVD) for 30 h compared to that of media control. mRNA levels were measured by qRT-PCR. **(B)** IL-8 protein concentration in the supernatants of 16HBE cells treated with iron chelators for 48 h. IL-8 was quantified by ELISA. Correlation between *IL8* gene expression and IL-8 protein production. All error bars represent SEM from three biological replicates.

**Fig. S8. Deferoxamine promotes the expression of proinflammatory genes in 16HBE CFTR mutants. (A)** Total RNA yield in wild-type 16HBE cells and 16HBE cells carrying mutations (G551D, ΔF508) in the cystic fibrosis transmembrane conductance regulator (CFTR) after 48 h treatment with 100 µM deferoxamine in serum-free EMEM. **(B)** Expression of proinflammatory genes (*NLRP3, IL1B, NLRP1, IL8, TNF*) or iron-regulated genes (*NDRG1*, *TFRC*) in 16HBE cells treated with deferoxamine compared to that of media control. mRNA levels were measured by qRT-PCR. All error bars represent SEM from three biological replicates.

## References

1. Kollef MH, Chastre J, Fagon JY, Francois B, Niederman MS, Rello J, Torres A, Vincent JL, Wunderink RG, Go KW, Rehm C. 2014. Global prospective epidemiologic and surveillance study of ventilator-associated pneumonia due to Pseudomonas aeruginosa. Crit Care Med 42:2178–87.

2. Lyczak JB, Cannon CL, Pier GB. 2002. Lung infections associated with cystic fibrosis. Clin Microbiol Rev 15:194–222.

3. Bhagirath AY, Li Y, Somayajula D, Dadashi M, Badr S, Duan K. 2016. Cystic fibrosis lung environment and Pseudomonas aeruginosa infection. BMC Pulm Med 16:174.

4. Hassett DJ, Borchers MT, Panos RJ. 2014. Chronic obstructive pulmonary disease (COPD): evaluation from clinical, immunological and bacterial pathogenesis perspectives. J Microbiol 52:211–26.

5. Murphy TF. 2009. Pseudomonas aeruginosa in adults with chronic obstructive pulmonary disease. Curr Opin Pulm Med 15:138–42.

6. Anderson GG, O’Toole GA. 2008. Innate and induced resistance mechanisms of bacterial biofilms. Curr Top Microbiol Immunol 322:85–105.

7. Moreau-Marquis S, Stanton BA, O’Toole GA. 2008. Pseudomonas aeruginosa biofilm formation in the cystic fibrosis airway. Pulm Pharmacol Ther 21:595–9.

8. Curran CS, Bolig T, Torabi-Parizi P. 2018. Mechanisms and Targeted Therapies for Pseudomonas aeruginosa Lung Infection. Am J Respir Crit Care Med 197:708–727.

9. Cézard C, Farvacques N, Sonnet P. 2015. Chemistry and biology of pyoverdines, Pseudomonas primary siderophores. Curr Med Chem 22:165–86.

10. Vasil ML, Ochsner UA. 1999. The response of Pseudomonas aeruginosa to iron: genetics, biochemistry and virulence. Mol Microbiol 34:399–413.

11. Cornelis P, Dingemans J. 2013. Pseudomonas aeruginosa adapts its iron uptake strategies in function of the type of infections. Front Cell Infect Microbiol 3:75.

12. Visca P, Imperi F, Lamont IL. 2007. Pyoverdine siderophores: from biogenesis to biosignificance. Trends Microbiol 15:22–30.

13. Xiao R, Kisaalita WS. 1997. Iron acquisition from transferrin and lactoferrin by Pseudomonas aeruginosa pyoverdin. Microbiology 143 (Pt 7):2509–15.

14. Dumas Z, Ross-Gillespie A, Kummerli R. 2013. Switching between apparently redundant iron-uptake mechanisms benefits bacteria in changeable environments. Proc Biol Sci 280:20131055.

15. Banin E, Vasil ML, Greenberg EP. 2005. Iron and Pseudomonas aeruginosa biofilm formation. Proc Natl Acad Sci U S A 102:11076–81.

16. Kang D, Kirienko NV. 2018. Interdependence between iron acquisition and biofilm formation in Pseudomonas aeruginosa. J Microbiol 56:449–457.

17. Minandri F, Imperi F, Frangipani E, Bonchi C, Visaggio D, Facchini M, Pasquali P, Bragonzi A, Visca P. 2016. Role of Iron Uptake Systems in Pseudomonas aeruginosa Virulence and Airway Infection. Infect Immun 84:2324–35.

18. Lamont IL, Beare PA, Ochsner U, Vasil AI, Vasil ML. 2002. Siderophore-mediated signaling regulates virulence factor production in Pseudomonasaeruginosa. Proc Natl Acad Sci U S A 99:7072–7.

19. Kirienko NV, Kirienko DR, Larkins-Ford J, Wählby C, Ruvkun G, Ausubel FM. 2013. Pseudomonas aeruginosa disrupts Caenorhabditis elegans iron homeostasis, causing a hypoxic response and death. Cell Host Microbe 13:406–16.

20. Kirienko NV, Ausubel FM, Ruvkun G. 2015. Mitophagy confers resistance to siderophore-mediated killing by Pseudomonas aeruginosa. Proc Natl Acad Sci U S A 112:1821–6.

21. Kang D, Kirienko DR, Webster P, Fisher AL, Kirienko NV. 2018. Pyoverdine, a siderophore from Pseudomonas aeruginosa, translocates into C. elegans, removes iron, and activates a distinct host response. Virulence 9:804–817.

22. Kang D, Revtovich AV, Chen Q, Shah KN, Cannon CL, Kirienko NV. 2019. Pyoverdine-Dependent Virulence of Pseudomonas aeruginosa Isolates From Cystic Fibrosis Patients. Front Microbiol 10:2048.

23. Imperi F, Massai F, Facchini M, Frangipani E, Visaggio D, Leoni L, Bragonzi A, Visca P. 2013. Repurposing the antimycotic drug flucytosine for suppression of Pseudomonas aeruginosa pathogenicity. Proc Natl Acad Sci U S A 110:7458–63.

24. Takase H, Nitanai H, Hoshino K, Otani T. 2000. Impact of siderophore production on Pseudomonas aeruginosa infections in immunosuppressed mice. Infect Immun 68:1834–9.

25. Meyer JM, Neely A, Stintzi A, Georges C, Holder IA. 1996. Pyoverdin is essential for virulence of Pseudomonas aeruginosa. Infect Immun 64:518–23.

26. Kang D, Kirienko NV. 2020. An In Vitro Cell Culture Model for Pyoverdine-Mediated Virulence. Pathogens 10.

27. Xu Q, Kang D, Meyer MD, Pennington CL, Gopal C, Schertzer JW, Kirienko NV. 2023. Cytotoxic rhamnolipid micelles drive acute virulence in Pseudomonas aeruginosa. bioRxiv doi:10.1101/2023.10.13.562257:2023.10.13.562257.

28. Wu Y, Yeh FL, Mao F, Chapman ER. 2009. Biophysical characterization of styryl dye-membrane interactions. Biophys J 97:101–9.

29. Jensen PO, Bjarnsholt T, Phipps R, Rasmussen TB, Calum H, Christoffersen L, Moser C, Williams P, Pressler T, Givskov M, Hoiby N. 2007. Rapid necrotic killing of polymorphonuclear leukocytes is caused by quorum-sensing-controlled production of rhamnolipid by Pseudomonas aeruginosa. Microbiology (Reading) 153:1329–1338.

30. Pearson JP, Pesci EC, Iglewski BH. 1997. Roles of Pseudomonas aeruginosa las and rhl quorum-sensing systems in control of elastase and rhamnolipid biosynthesis genes. J Bacteriol 179:5756–67.

31. Winson MK, Swift S, Fish L, Throup JP, Jorgensen F, Chhabra SR, Bycroft BW, Williams P, Stewart GS. 1998. Construction and analysis of luxCDABE-based plasmid sensors for investigating N-acyl homoserine lactone-mediated quorum sensing. FEMS Microbiol Lett 163:185–92.

32. Lee J, Zhang L. 2015. The hierarchy quorum sensing network in Pseudomonas aeruginosa. Protein Cell 6:26–41.

33. Filloux A. 2011. Protein Secretion Systems in Pseudomonas aeruginosa: An Essay on Diversity, Evolution, and Function. Front Microbiol 2:155.

34. Kohler T, Epp SF, Curty LK, Pechere JC. 1999. Characterization of MexT, the regulator of the MexE-MexF-OprN multidrug efflux system of Pseudomonas aeruginosa. J Bacteriol 181:6300–5.

35. Kohler T, van Delden C, Curty LK, Hamzehpour MM, Pechere JC. 2001. Overexpression of the MexEF-OprN multidrug efflux system affects cell-to-cell signaling in Pseudomonas aeruginosa. J Bacteriol 183:5213–22.

36. Vaillancourt M, Limsuwannarot SP, Bresee C, Poopalarajah R, Jorth P. 2021. Pseudomonas aeruginosa mexR and mexEF Antibiotic Efflux Pump Variants Exhibit Increased Virulence. Antibiotics (Basel) 10.

37. Kirienko DR, Revtovich AV, Kirienko NV. 2016. A High-Content, Phenotypic Screen Identifies Fluorouridine as an Inhibitor of Pyoverdine Biosynthesis and Pseudomonas aeruginosa Virulence. mSphere 1.

38. Kang D, Revtovich AV, Deyanov AE, Kirienko NV. 2021. Pyoverdine Inhibitors and Gallium Nitrate Synergistically Affect Pseudomonas aeruginosa. mSphere doi:10.1128/mSphere.00401-21:e0040121.

39. Costabile G, d’Angelo I, d’Emmanuele di Villa Bianca R, Mitidieri E, Pompili B, Del Porto P, Leoni L, Visca P, Miro A, Quaglia F, Imperi F, Sorrentino R, Ungaro F. 2016. Development of inhalable hyaluronan/mannitol composite dry powders for flucytosine repositioning in local therapy of lung infections. J Control Release 238:80–91.

40. Oliver JM, Berlin RD, Davis BH. 1984. Use of horseradish peroxidase and fluorescent dextrans to study fluid pinocytosis in leukocytes. Methods Enzymol 108:336–47.

41. Holden VI, Lenio S, Kuick R, Ramakrishnan SK, Shah YM, Bachman MA. 2014. Bacterial siderophores that evade or overwhelm lipocalin 2 induce hypoxia inducible factor 1α and proinflammatory cytokine secretion in cultured respiratory epithelial cells. Infect Immun 82:3826–36.

42. Lee HJ, Lee J, Lee SK, Lee SK, Kim EC. 2007. Differential regulation of iron chelator-induced IL-8 synthesis via MAP kinase and NF-kappaB in immortalized and malignant oral keratinocytes. BMC Cancer 7:176.

43. Choi EY, Kim EC, Oh HM, Kim S, Lee HJ, Cho EY, Yoon KH, Kim EA, Han WC, Choi SC, Hwang JY, Park C, Oh BS, Kim Y, Kimm KC, Park KI, Chung HT, Jun CD. 2004. Iron chelator triggers inflammatory signals in human intestinal epithelial cells: involvement of p38 and extracellular signal-regulated kinase signaling pathways. J Immunol 172:7069–77.

44. Moldoveanu B, Otmishi P, Jani P, Walker J, Sarmiento X, Guardiola J, Saad M, Yu J. 2009. Inflammatory mechanisms in the lung. J Inflamm Res 2:1–11.

45. Hauser AR. 2009. The type III secretion system of Pseudomonas aeruginosa: infection by injection. Nat Rev Microbiol 7:654–65.

46. Michalska M, Wolf P. 2015. Pseudomonas Exotoxin A: optimized by evolution for effective killing. Front Microbiol 6:963.

47. Hall S, McDermott C, Anoopkumar-Dukie S, McFarland AJ, Forbes A, Perkins AV, Davey AK, Chess-Williams R, Kiefel MJ, Arora D, Grant GD. 2016. Cellular Effects of Pyocyanin, a Secreted Virulence Factor of Pseudomonas aeruginosa. Toxins (Basel) 8.

48. Thi MTT, Wibowo D, Rehm BHA. 2020. Pseudomonas aeruginosa Biofilms. Int J Mol Sci 21.

49. Ochsner UA, Johnson Z, Lamont IL, Cunliffe HE, Vasil ML. 1996. Exotoxin A production in Pseudomonas aeruginosa requires the iron-regulated pvdS gene encoding an alternative sigma factor. Mol Microbiol 21:1019–28.

50. Plotkowski MC, Povoa HC, Zahm JM, Lizard G, Pereira GM, Tournier JM, Puchelle E. 2002. Early mitochondrial dysfunction, superoxide anion production, and DNA degradation are associated with non-apoptotic death of human airway epithelial cells induced by Pseudomonas aeruginosa exotoxin A. Am J Respir Cell Mol Biol 26:617–26.

51. Azghani AO. 1996. Pseudomonas aeruginosa and epithelial permeability: role of virulence factors elastase and exotoxin A. Am J Respir Cell Mol Biol 15:132–40.

52. Hirakata Y, Furuya N, Tateda K, Kaku M, Yamaguchi K. 1993. In vivo production of exotoxin A and its role in endogenous Pseudomonas aeruginosa septicemia in mice. Infect Immun 61:2468–73.

53. Pillar CM, Hobden JA. 2002. Pseudomonas aeruginosa exotoxin A and keratitis in mice. Invest Ophthalmol Vis Sci 43:1437–44.

54. Miyazaki S, Matsumoto T, Tateda K, Ohno A, Yamaguchi K. 1995. Role of exotoxin A in inducing severe Pseudomonas aeruginosa infections in mice. J Med Microbiol 43:169–75.

55. Malloy JL, Veldhuizen RA, Thibodeaux BA, O’Callaghan RJ, Wright JR. 2005. Pseudomonas aeruginosa protease IV degrades surfactant proteins and inhibits surfactant host defense and biophysical functions. Am J Physiol Lung Cell Mol Physiol 288:L409–18.

56. Bradshaw JL, Caballero AR, Bierdeman MA, Adams KV, Pipkins HR, Tang A, O’Callaghan RJ, McDaniel LS. 2018. Pseudomonas aeruginosa Protease IV Exacerbates Pneumococcal Pneumonia and Systemic Disease. mSphere 3.

57. Guillon A, Brea D, Morello E, Tang A, Jouan Y, Ramphal R, Korkmaz B, Perez-Cruz M, Trottein F, O’Callaghan RJ, Gosset P, Si-Tahar M. 2017. Pseudomonas aeruginosa proteolytically alters the interleukin 22-dependent lung mucosal defense. Virulence 8:810–820.

58. Engel LS, Hobden JA, Moreau JM, Callegan MC, Hill JM, O’Callaghan RJ. 1997. Pseudomonas deficient in protease IV has significantly reduced corneal virulence. Invest Ophthalmol Vis Sci 38:1535–42.

59. Engel LS, Hill JM, Moreau JM, Green LC, Hobden JA, O’Callaghan RJ. 1998. Pseudomonas aeruginosa protease IV produces corneal damage and contributes to bacterial virulence. Invest Ophthalmol Vis Sci 39:662–5.

60. Gdaniec BG, Bonini F, Prodon F, Braschler T, Kohler T, van Delden C. 2022. Pseudomonas aeruginosa rhamnolipid micelles deliver toxic metabolites and antibiotics into Staphylococcus aureus. iScience 25:103669.

61. McClure CD, Schiller NL. 1992. Effects of Pseudomonas aeruginosa rhamnolipids on human monocyte-derived macrophages. J Leukoc Biol 51:97–102.

62. Van Gennip M, Christensen LD, Alhede M, Phipps R, Jensen PO, Christophersen L, Pamp SJ, Moser C, Mikkelsen PJ, Koh AY, Tolker-Nielsen T, Pier GB, Hoiby N, Givskov M, Bjarnsholt T. 2009. Inactivation of the rhlA gene in Pseudomonas aeruginosa prevents rhamnolipid production, disabling the protection against polymorphonuclear leukocytes. APMIS 117:537–46.

63. Zulianello L, Canard C, Kohler T, Caille D, Lacroix JS, Meda P. 2006. Rhamnolipids are virulence factors that promote early infiltration of primary human airway epithelia by Pseudomonas aeruginosa. Infect Immun 74:3134–47.

64. Read RC, Roberts P, Munro N, Rutman A, Hastie A, Shryock T, Hall R, McDonald-Gibson W, Lund V, Taylor G, et al. 1992. Effect of Pseudomonas aeruginosa rhamnolipids on mucociliary transport and ciliary beating. J Appl Physiol (1985) 72:2271–7.

65. Hentzer M, Wu H, Andersen JB, Riedel K, Rasmussen TB, Bagge N, Kumar N, Schembri MA, Song Z, Kristoffersen P, Manefield M, Costerton JW, Molin S, Eberl L, Steinberg P, Kjelleberg S, Høiby N, Givskov M. 2003. Attenuation of Pseudomonas aeruginosa virulence by quorum sensing inhibitors. EMBO J 22:3803–15.

66. Kang D, Turner KE, Kirienko NV. 2017. PqsA Promotes Pyoverdine Production via Biofilm Formation. Pathogens 7.

67. Stintzi A, Evans K, Meyer JM, Poole K. 1998. Quorum-sensing and siderophore biosynthesis in Pseudomonas aeruginosa: lasR/lasI mutants exhibit reduced pyoverdine biosynthesis. FEMS Microbiol Lett 166:341–5.

68. Thomann A, de Mello Martins AG, Brengel C, Empting M, Hartmann RW. 2016. Application of Dual Inhibition Concept within Looped Autoregulatory Systems toward Antivirulence Agents against Pseudomonas aeruginosa Infections. ACS Chem Biol 11:1279–86.

69. R YR-R, Salvador MJ. 2020. Phenotypic detection of quorum sensing inhibition in Pseudomonas aeruginosa pyoverdine and swarming by volatile organic products. Future Microbiol 15:1147–1156.

70. Holden VI, Breen P, Houle S, Dozois CM, Bachman MA. 2016. Klebsiella pneumoniae Siderophores Induce Inflammation, Bacterial Dissemination, and HIF-1α Stabilization during Pneumonia. MBio 7.

71. Marvig RL, Damkiær S, Khademi SM, Markussen TM, Molin S, Jelsbak L. 2014. Within-host evolution of Pseudomonas aeruginosa reveals adaptation toward iron acquisition from hemoglobin. MBio 5:e00966–14.

72. De Vos D, De Chial M, Cochez C, Jansen S, Tummler B, Meyer JM, Cornelis P. 2001. Study of pyoverdine type and production by Pseudomonas aeruginosa isolated from cystic fibrosis patients: prevalence of type II pyoverdine isolates and accumulation of pyoverdine-negative mutations. Arch Microbiol 175:384–8.

73. Andersen SB, Marvig RL, Molin S, Krogh Johansen H, Griffin AS. 2015. Long-term social dynamics drive loss of function in pathogenic bacteria. Proc Natl Acad Sci U S A 112:10756–61.

74. Martin LW, Reid DW, Sharples KJ, Lamont IL. 2011. Pseudomonas siderophores in the sputum of patients with cystic fibrosis. Biometals 24:1059–67.

75. Haas B, Murphy E, Castignetti D. 1991. Siderophore synthesis by mucoid Pseudomonas aeruginosa strains isolated from cystic fibrosis patients. Can J Microbiol 37:654–7.

76. Mayer-Hamblett N, Rosenfeld M, Gibson RL, Ramsey BW, Kulasekara HD, Retsch-Bogart GZ, Morgan W, Wolter DJ, Pope CE, Houston LS, Kulasekara BR, Khan U, Burns JL, Miller SI, Hoffman LR. 2014. Pseudomonas aeruginosa in vitro phenotypes distinguish cystic fibrosis infection stages and outcomes. Am J Respir Crit Care Med 190:289–97.

77. Kang D, Zhang L, Kirienko NV. 2021. High-Throughput Approaches for the Identification of Pseudomonas aeruginosa Antivirulents. mBio 12.

78. Kaneko Y, Thoendel M, Olakanmi O, Britigan BE, Singh PK. 2007. The transition metal gallium disrupts Pseudomonas aeruginosa iron metabolism and has antimicrobial and antibiofilm activity. The Journal of Clinical Investigation 117:877–888.

79. Goss CH, Kaneko Y, Khuu L, Anderson GD, Ravishankar S, Aitken ML, Lechtzin N, Zhou G, Czyz DM, McLean K, Olakanmi O, Shuman HA, Teresi M, Wilhelm E, Caldwell E, Salipante SJ, Hornick DB, Siehnel RJ, Becker L, Britigan BE, Singh PK. 2018. Gallium disrupts bacterial iron metabolism and has therapeutic effects in mice and humans with lung infections. Sci Transl Med 10.

80. Britigan BE, Rasmussen GT, Cox CD. 1994. Pseudomonas siderophore pyochelin enhances neutrophil-mediated endothelial cell injury. Am J Physiol 266:L192–8.

81. Ong KS, Cheow YL, Lee SM. 2017. The role of reactive oxygen species in the antimicrobial activity of pyochelin. J Adv Res 8:393–398.

82. Adler C, Corbalan NS, Seyedsayamdost MR, Pomares MF, de Cristobal RE, Clardy J, Kolter R, Vincent PA. 2012. Catecholate siderophores protect bacteria from pyochelin toxicity. PLoS One 7:e46754.

83. Jenul C, Keim KC, Jens JN, Zeiler MJ, Schilcher K, Schurr MJ, Melander C, Phelan VV, Horswill AR. 2023. Pyochelin biotransformation by Staphylococcusaureus shapes bacterial competition with Pseudomonas aeruginosa in polymicrobial infections. Cell Rep 42:112540.

84. Kruger P, Saffarzadeh M, Weber AN, Rieber N, Radsak M, von Bernuth H, Benarafa C, Roos D, Skokowa J, Hartl D. 2015. Neutrophils: Between host defence, immune modulation, and tissue injury. PLoS Pathog 11:e1004651.

85. Twigg MS, Brockbank S, Lowry P, FitzGerald SP, Taggart C, Weldon S. 2015. The Role of Serine Proteases and Antiproteases in the Cystic Fibrosis Lung. Mediators Inflamm 2015:293053.

86. Cantin AM, Hartl D, Konstan MW, Chmiel JF. 2015. Inflammation in cystic fibrosis lung disease: Pathogenesis and therapy. J Cyst Fibros 14:419–30.

87. Hosseinian N, Cho Y, Lockey RF, Kolliputi N. 2015. The role of the NLRP3 inflammasome in pulmonary diseases. Ther Adv Respir Dis 9:188–97.

88. McElvaney OJ, Zaslona Z, Becker-Flegler K, Palsson-McDermott EM, Boland F, Gunaratnam C, Gulbins E, O’Neill LA, Reeves EP, McElvaney NG. 2019. Specific Inhibition of the NLRP3 Inflammasome as an Antiinflammatory Strategy in Cystic Fibrosis. Am J Respir Crit Care Med 200:1381–1391.

89. McKee CM, Coll RC. 2020. NLRP3 inflammasome priming: A riddle wrapped in a mystery inside an enigma. J Leukoc Biol 108:937–952.

90. Estivill X, Bancells C, Ramos C. 1997. Geographic distribution and regional origin of 272 cystic fibrosis mutations in European populations. The Biomed CF Mutation Analysis Consortium. Hum Mutat 10:135–54.

91. de Albuquerque Wanderley Sales V, Timoteo TRR, da Silva NM, de Melo CG, Ferreira AS, de Oliveira MVG, de Oliveira Silva E, Dos Santos Mendes LM, Rolim LA, Neto PJR. 2021. A Systematic Review of the Anti-inflammatory Effects of Gallium Compounds. Curr Med Chem 28:2062–2076.

92. Zhang C, Yang B, Biazik JM, Webster RF, Xie W, Tang J, Allioux FM, Abbasi R, Mousavi M, Goldys EM, Kilian KA, Chandrawati R, Esrafilzadeh D, Kalantar-Zadeh K. 2022. Gallium Nanodroplets are Anti-Inflammatory without Interfering with Iron Homeostasis. ACS Nano 16:8891–8903.

93. Apseloff G. 1999. Therapeutic uses of gallium nitrate: past, present, and future. Am J Ther 6:327–39.

94. Jacobs MA, Alwood A, Thaipisuttikul I, Spencer D, Haugen E, Ernst S, Will O, Kaul R, Raymond C, Levy R, Chun-Rong L, Guenthner D, Bovee D, Olson MV, Manoil C. 2003. Comprehensive transposon mutant library of Pseudomonas aeruginosa. Proc Natl Acad Sci U S A 100:14339–44.

95. Liberati NT, Urbach JM, Miyata S, Lee DG, Drenkard E, Wu G, Villanueva J, Wei T, Ausubel FM. 2006. An ordered, nonredundant library of Pseudomonas aeruginosa strain PA14 transposon insertion mutants. Proc Natl Acad Sci U S A 103:2833–8.

96. Bankevich A, Nurk S, Antipov D, Gurevich AA, Dvorkin M, Kulikov AS, Lesin VM, Nikolenko SI, Pham S, Prjibelski AD, Pyshkin AV, Sirotkin AV, Vyahhi N, Tesler G, Alekseyev MA, Pevzner PA. 2012. SPAdes: a new genome assembly algorithm and its applications to single-cell sequencing. J Comput Biol 19:455–77.

97. Seemann T. 2014. Prokka: rapid prokaryotic genome annotation. Bioinformatics 30:2068–9.

98. Deatherage DE, Barrick JE. 2014. Identification of mutations in laboratory-evolved microbes from next-generation sequencing data using breseq. Methods Mol Biol 1151:165–88.

