## Supplementary figures and images for "*In vitro* Lung Epithelial Cell Model Reveals Novel Roles for *Pseudomonas aeruginosa* Siderophores"

### Supplemental Figures 1-8

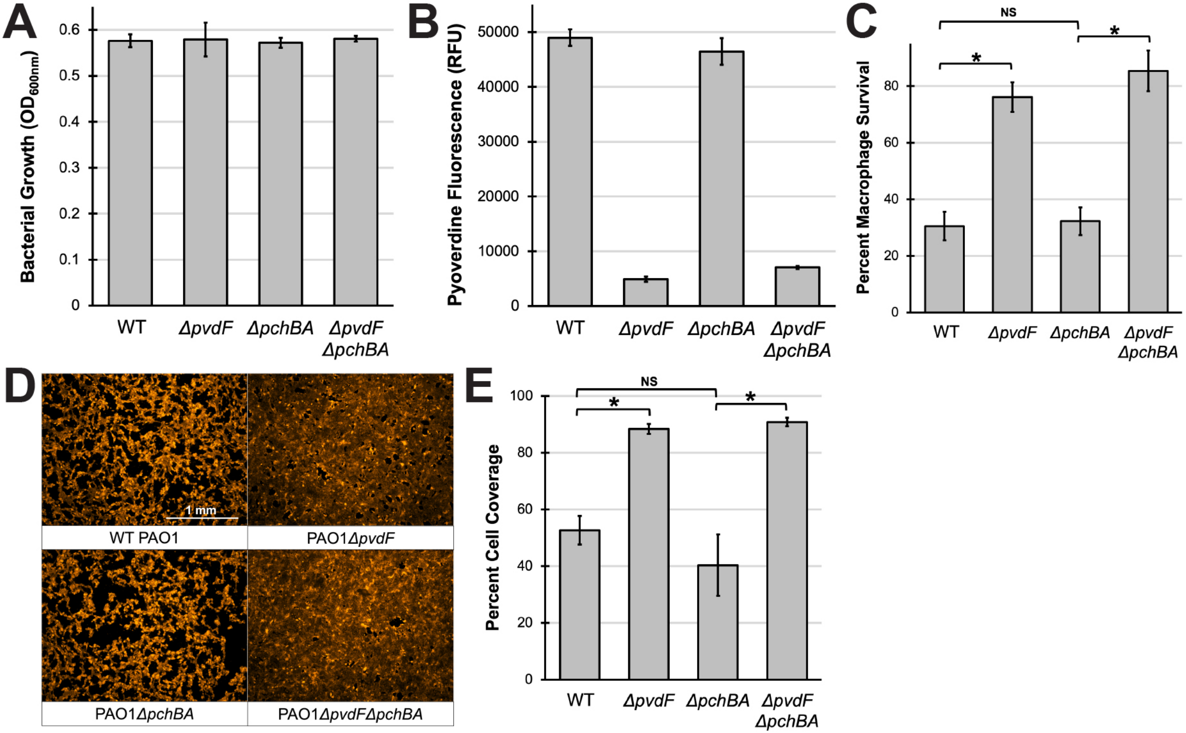

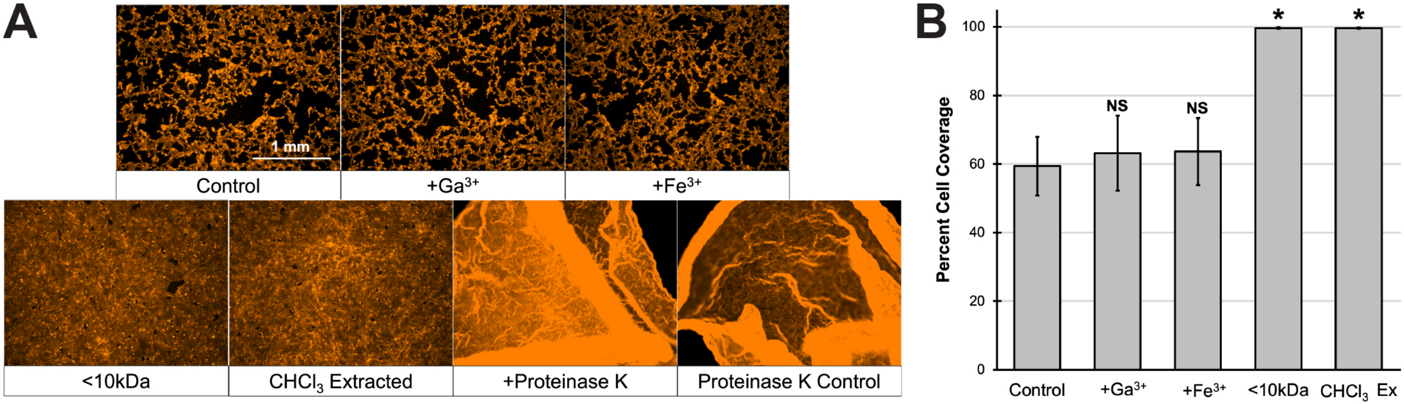

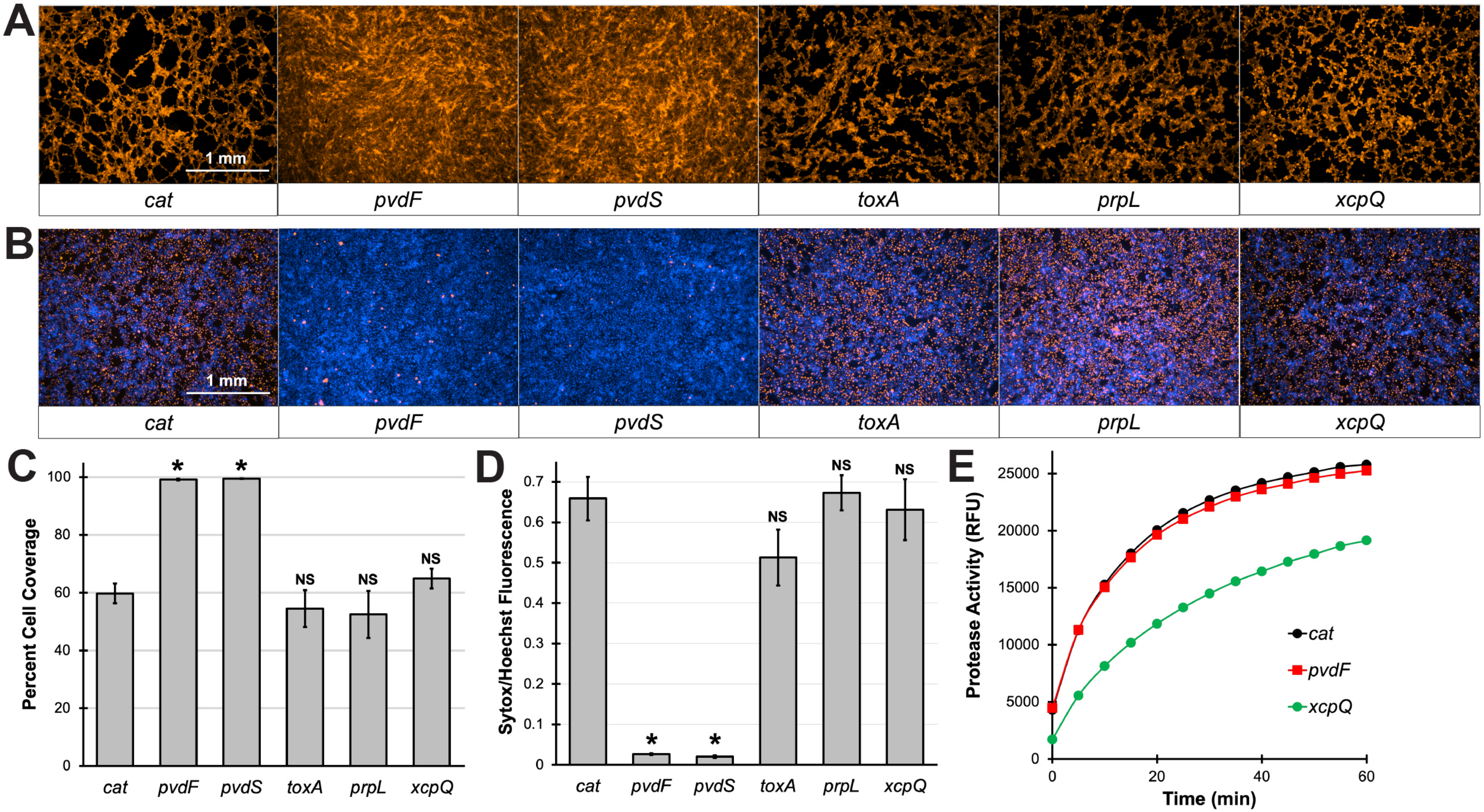

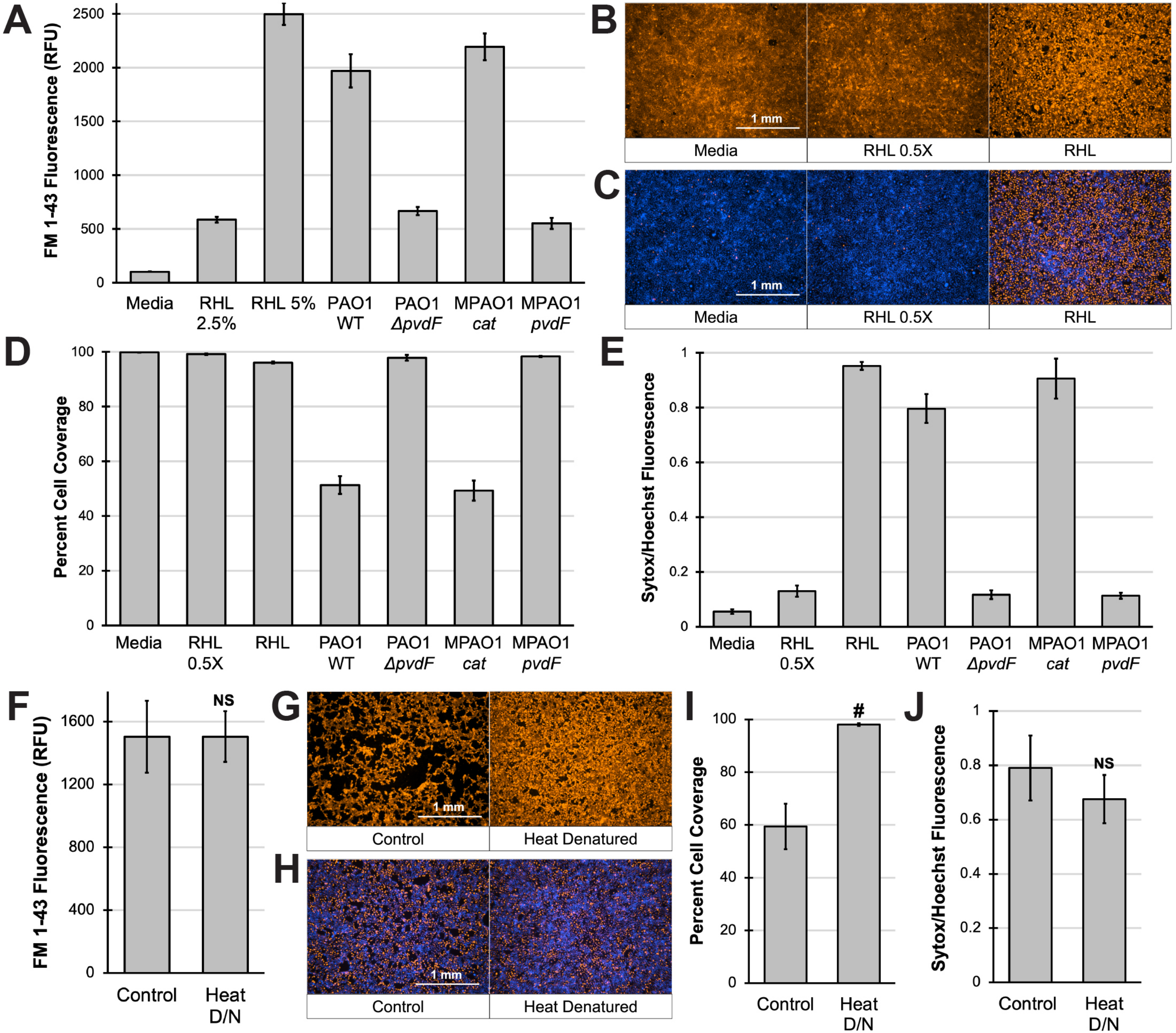

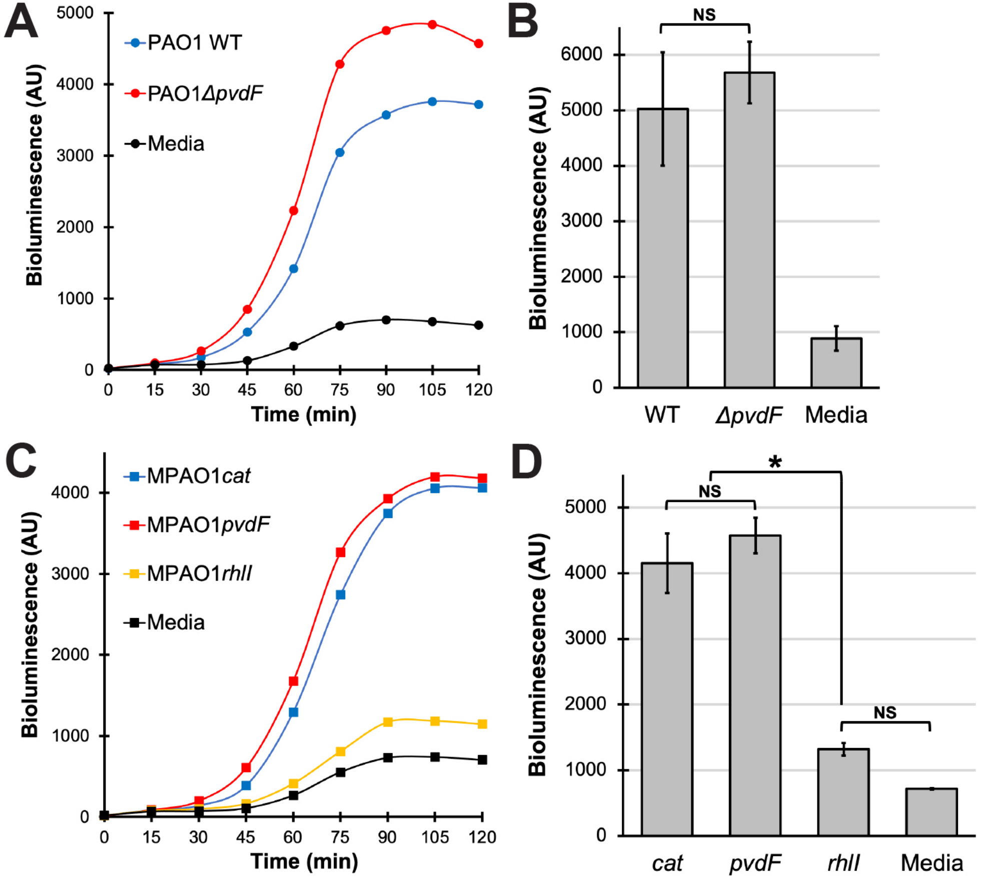

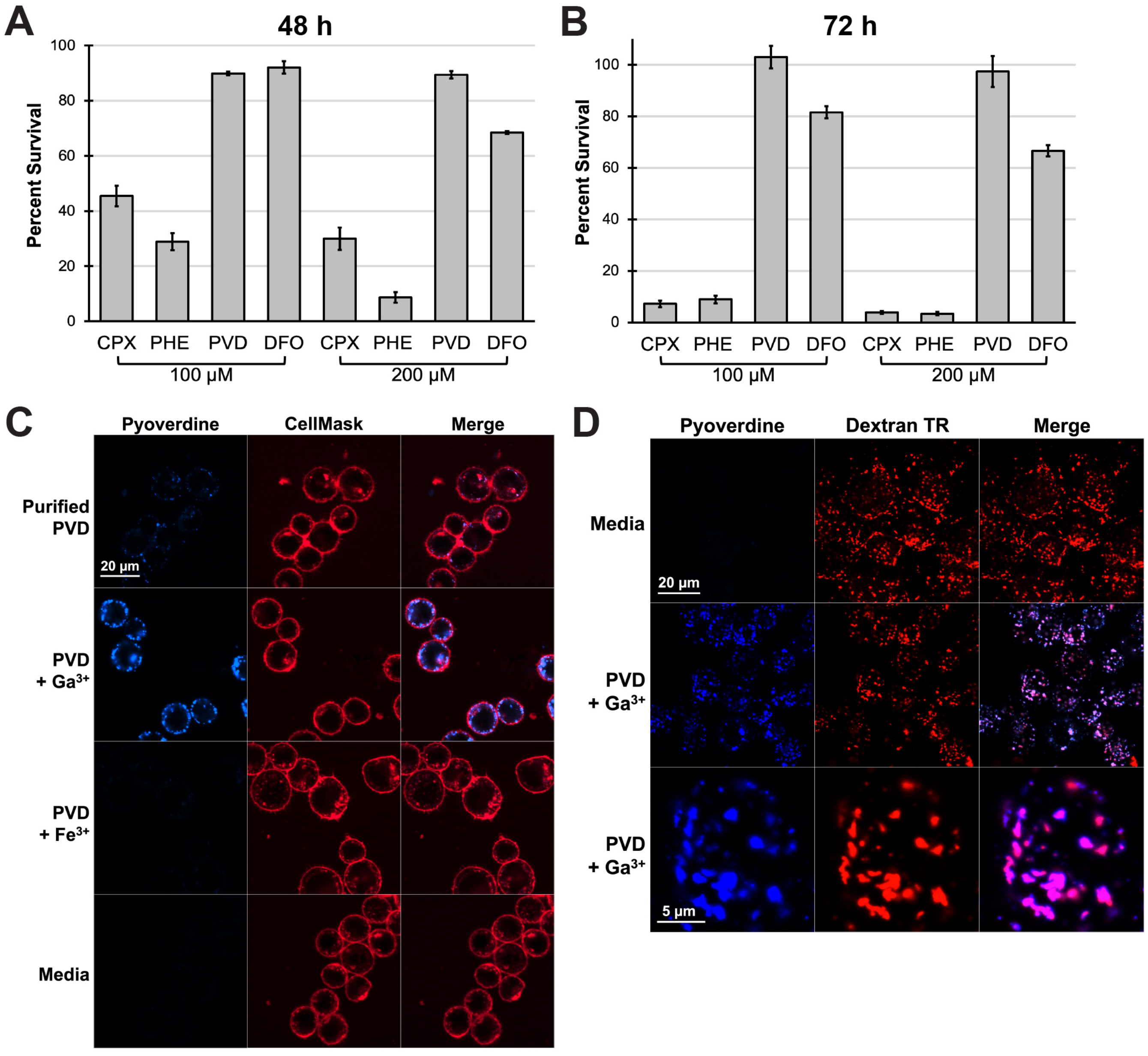

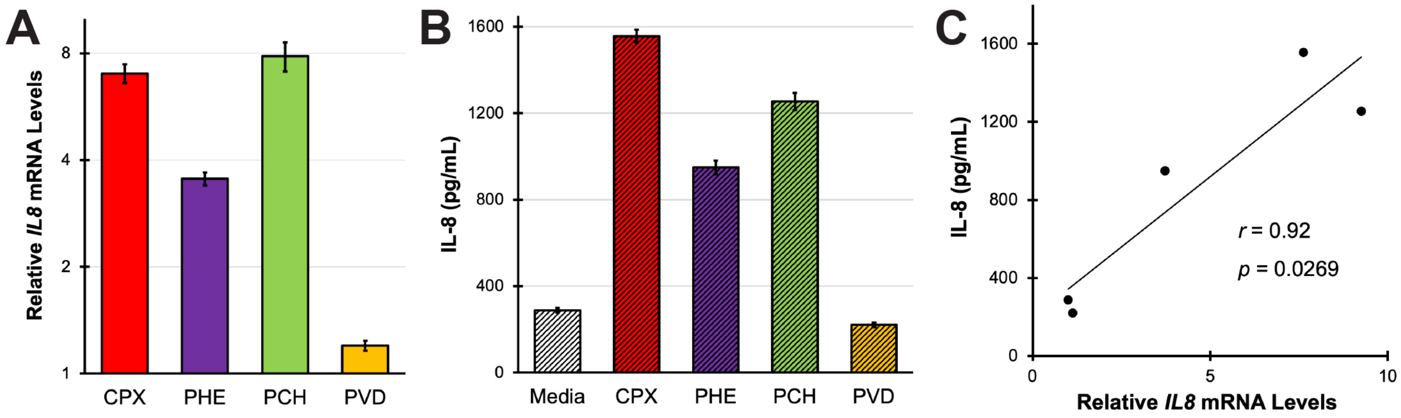

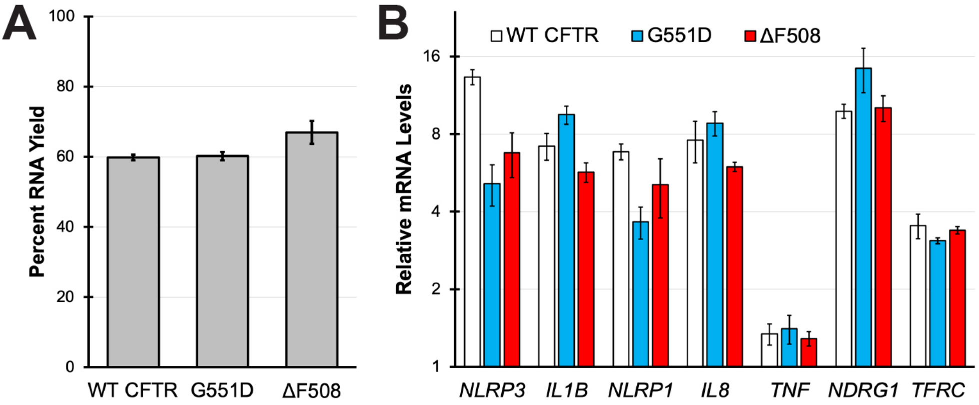
