## Supplemental Table S1 for "*In vitro* Lung Epithelial Cell Model Reveals Novel Roles for *Pseudomonas aeruginosa* Siderophores"

**Table S1. List of primers used in this study.**

| Primer Name | Sequence (5' - 3') | Description |
|---|---|---|
| ARB1D | GGCCAGGCCTGCAGATGATGNNNNNNNNNNGTAT | MPAO1 Tn Verification Arbitrary PCR Round 1 |
| PhoA Tn_1 | GTTAACCATAACTTCGTATAATG | MPAO1 Tn Verification Arbitrary PCR Round 1 (ISphoA/hah) |
| LacZ Tn_1 | GTAAACTGGATGGCTTTCTTGCC | MPAO1 Tn Verification Arbitrary PCR Round 1 (ISlacZ/hah) |
| ARB2A | GGCCAGGCCTGCAGATGATG | MPAO1 Tn Verification Arbitrary PCR Round 2 |
| PhoA Tn_2 | AATTGGATAACTTCGTATAATGTATGC | MPAO1 Tn Verification Arbitrary PCR Round 2 (ISphoA/hah) |
| LacZ Tn_2 | AAGGATCTGATGGCGCAGGGGATCCCC | MPAO1 Tn Verification Arbitrary PCR Round 2 (ISlacZ/hah) |
| MPAO1_Seq | GTTATTAATTAAGCATCACC | MPAO1 Tn Verification Sanger Sequencing Primer |
| mexT-F1 | TATGGGCGGGGAAACTGGCCACGCG | *P. aeruginosa mexT* Full Gene PCR Round 1 -Forward |
| mexT-R1 | CTTTCTTCACCAGTGCGCCTTCATG | *P. aeruginosa mexT* Full Gene PCR Round 1 - Reverse |
| mexT-F2 | AGCTCTTCGCATTTGAGGACCTCTG | *P. aeruginosa mexT* Full Gene PCR Round 2 -Forward |
| mexT-R2 | AGCGCCAGGAGAAGTGGGATGACTG | *P. aeruginosa mexT* Full Gene PCR Round 2 - Reverse |
| mexT-Seq | TGCCTGTCAGTGATCCTATG | *P. aeruginosa mexT* Sanger Sequencing Primer |
| gyrB_F | CCTGCTGTTGACCTTCTTCT | *P. aeruginosa gyrB* qRT-PCR - Forward |
| gyrB_R | CTGGTCGTCCTTGATGTACTG | *P. aeruginosa gyrB* qRT-PCR - Reverse |
| mexE_F | CGGCAACCTGGTCAACT | *P. aeruginosa mexE* qRT-PCR - Forward |
| mexE_R | CTCGACGTACTTGAGGAACAC | *P. aeruginosa mexE* qRT-PCR - Reverse |
| mexF_F | GAAGCTCCCGGAAGAAGTG | *P. aeruginosa mexF* qRT-PCR - Forward |
| mexF_R | AGCATGTCGTAGCGGTTATC | *P. aeruginosa mexF* qRT-PCR - Reverse |
| ACTB_F | CACCATTGGCAATGAGCGGTTC | 16HBE *ACTB* qRT-PCR - Forward |
| ACTB_R | AGGTCTTTGCGGATGTCCACGT | 16HBE *ACTB* qRT-PCR - Reverse |
| NLRP3_F | GGACTGAAGCACCTGTTGTGCA | 16HBE *NLRP3* qRT-PCR - Forward |
| NLRP3_R | TCCTGAGTCTCCCAAGGCATTC | 16HBE *NLRP3* qRT-PCR - Reverse |
| IL1B_F | CCACAGACCTTCCAGGAGAATG | 16HBE *IL1B* qRT-PCR - Forward |
| IL1B_R | GTGCAGTTCAGTGATCGTACAGG | 16HBE *IL1B* qRT-PCR - Reverse |
| NRLP1_F | ATTGAGGGCAGGCAGCACAGAT | 16HBE *NLRP1* qRT-PCR - Forward |
| NRLP1_R | CTCCTTCAGGTTTCTGGTGACC | 16HBE *NLRP1* qRT-PCR - Reverse |
| IL8_F | GAGAGTGATTGAGAGTGGACCAC | 16HBE *IL8* qRT-PCR - Forward |
| IL8_R | CACAACCCTCTGCACCCAGTTT | 16HBE *IL8* qRT-PCR - Reverse |
| TNF_F | CTCTTCTGCCTGCTGCACTTTG | 16HBE *TNF* qRT-PCR - Forward |
| TNF_R | ATGGGCTACAGGCTTGTCACTC | 16HBE *TNF* qRT-PCR - Reverse |
| NDRG1_F | ATCACCCAGCACTTTGCCGTCT | 16HBE *NDRG1* qRT-PCR - Forward |
| NDRG1_R | GACTCCAGGAAGCATTTCAGCC | 16HBE *NDRG1* qRT-PCR - Reverse |
| TFRC_F | ATCGGTTGGTGCCACTGAATGG | 16HBE *TFRC* qRT-PCR - Forward |
| TFRC_R | ACAACAGTGGGCTGGCAGAAAC | 16HBE *TFRC* qRT-PCR - Reverse |
